# Targeting Neuropilin-1 to Enhance Immunotherapy in Melanoma: Reducing Peripheral Treg-Mediated Immunosuppression and Tumour Progression

**DOI:** 10.1101/2024.09.27.615359

**Authors:** Somlata Khamaru, Kshyama Subhadarsini Tung, Subhasis Chattopadhyay

**Affiliations:** National Institute of Science Education and Research, School of Biological Sciences, Bhubaneswar, Odisha 752050, Odisha, India; Homi Bhabha National Institute, Training School Complex, Anushaktinagar, Mumbai, 400094, India

## Abstract

Melanoma, the most aggressive type of skin cancer with a high mutation rate, is the fifth most common cancer among Caucasians. Despite advancements in treatments like immune checkpoint inhibitors and targeted therapies, over 40% of patients experience immune-related side effects, presenting significant challenges. Neuropilin-1 (NRP1) has become an essential target in cancer therapy due to its overexpression in various cancers, where it enhances regulatory T cell (Treg) function and supports tumor growth, often leading to poor outcomes.

This study investigated the effects of NRP1 inhibition in B16-F10 melanoma and its impact on immune responses regulated by Tregs. NRP1 was overexpressed in several cancers, including B16-F10 cells, compared to non-cancerous NIH-3T3 cells. Inhibiting NRP1 selectively caused apoptosis in B16-F10 cells without affecting NIH-3T3 cells. It also reversed the immunosuppression of splenic T cells induced by B16-F10-conditioned media, reducing Treg markers (NRP1, NKG2A, FOXP3), Treg activity, and the production of immunosuppressive cytokines (IL-10, IL-17A). Furthermore, NRP1 inhibition increased T cell proliferation and boosted the release of effector cytokines (TNF, IFN-γ, IL-6, IL-2). NRP1 inhibition also suppressed the STAT, ERK MAPK, and Smad2/3 pathways while activating the PI3K/AKT pathway. In splenic T cells from B16-F10 tumor-bearing mice treated with an NRP1 inhibitor, there was a decrease in Treg markers and activity, along with enhanced T cell proliferation. Additionally, NRP1 inhibitor treatment reduced lung metastasis, decreased tumor size, and improved survival in these mice.

This study shows that inhibiting NRP1 may slow B16-F10 melanoma progression and reduce Treg-mediated immunosuppression. This suggests its potential as a promising approach in future cancer immunotherapies, especially in combination with other treatments.

## Introduction

Melanoma is a deadly skin cancer caused by the uncontrolled growth of melanocytes. Its incidence has risen sharply over the past 50 years, especially among Caucasians, leading to significant health and economic impacts due to the high loss of life years (1).

Melanoma, an immunogenic tumour, has a worse prognosis in immunosuppressive environments. Studies have identified several mechanisms of melanoma-induced immunosuppression, including down-regulation of surface antigens, mutations, reduced costimulatory signals, recruitment of immunosuppressive cells, secretion of immunosuppressive cytokines, and tolerance induction. Previous research also linked regulatory T cells (Tregs) (CD4^+^CD25^+^ FOXP3^+^) to poor responses to immunotherapy and increased numbers in advanced melanoma (2–4). In melanoma, increased levels of programmed cell death 1 protein (PD-1) in CD8^+^ T cells have been associated with higher apoptosis rates in these cells, weakening their ability to fight tumours (5). PD-1 on cytotoxic T cells interacts with programmed death ligand 1 (PD-L1) on tumour cells and immune cells like dendritic cells (DCs) and other antigen-presenting cells (APCs), leading to T-cell exhaustion (6). Other immune checkpoints, such as Neuropilin-1 (NRP1), cytotoxic T-lymphocyte associated protein 4 (CTLA-4), T-cell immunoglobulin and mucin-domain containing-3 (TIM-3), lymphocyte-activation gene 3 (LAG-3) and V-domain immunoglobulin suppressor of T cell activation (VISTA), also suppress T-cell activity, promoting tumour growth and metastasis (7–18). Regulatory T cells (Tregs) in the melanoma environment modulate the immune response by releasing cytokines like (interleukin-10 (IL-10), interleukin-35 (IL-35), transforming growth factor beta (TGF-β)), cytotoxicity against T cells, preventing DC maturation, and disrupting cell metabolism (16,19). Melanoma also impairs natural killer (NK) cell responses by expressing molecules like Fas, apoptosis antigen 1 (APO1), and Natural Killer Cell Receptor (NKG2A) ligands, which trigger apoptosis in immune cells (2). Additionally, melanoma cells secrete factors like vascular endothelial growth factor (VEGF), TGF-β, essential fibroblast growth factor (bFGF), galectins, Th2 cytokines, chemokines, indoleamine 2,3-dioxygenase (IDO), and exosomes, which promote immune tolerance and suppression (20).

Immune checkpoint inhibitors (ICIs), like anti-CTLA-4 and anti-PD-1 antibodies, have been shown to restore CD8^+^ T cell function, leading to significant tumour regression and long-term control in about 50% of advanced melanoma patients, compared to less than 10% in the past (12). However, most patients do not experience lasting benefits, even with combination ICIs (21). Research suggests this resistance is influenced by genomic factors, immune cell behaviour, the tumour microenvironment, host characteristics, and the gut microbiome (22). ICIs can cause immune-related adverse events (irAEs) and are expensive, making it essential to use them selectively. Identifying biomarkers to predict treatment response and adverse effects is crucial for improving cancer therapy (11,23). Checkpoint blockade therapy has been found to reverse T cell inhibition through receptors like LAG-3 and TIM-3, emphasising the need to balance effector T cell activation and Treg cell suppression to optimise clinical outcomes. Tailored therapies based on this balance may enhance treatment success in specific cancers and individual patients (23).

Neuropilin is a type I transmembrane protein with two homologues, Neuropilin-1 (NRP1) and Neuropilin-2 (NRP2), that functions as a receptor for class 3 semaphorins (e.g., Sema3A) and specific VEGF isoforms (24). Recently, NRP1 has attracted attention in immune-oncology for its role in enhancing Treg cell suppression and limiting durable effector T cell responses. Initially recognised as a Treg cell marker, NRP1 stabilises Tregs in the tumour microenvironment (TME), inhibiting CD8^+^ T cell anti-tumour activity. In cancer, NRP1 on Tregs promotes immune suppression by facilitating Treg recruitment through VEGF co-receptor activity and maintaining tumour-specific Treg stability via Sema4a binding, which modulates the Akt-mTOR signalling pathway and the PTEN-Akt-FoxO axis (25,26). Increased NRP1^+^ Tregs highlight its critical role in immune regulation within tumours. Additionally, NRP1 has emerged as a therapeutic target in malignant melanoma to reduce invasiveness, combat BRAF inhibitor resistance, inhibit Treg recruitment, and enhance immunotherapy effectiveness (27). Combining anti-NRP1 with anti-PD-1 immunotherapy has improved CD8^+^ T cell proliferation, cytotoxicity, and tumour control, suggesting that targeting NRP1 may be a promising immunotherapeutic strategy (28).

Inhibition of NRP1 by depletion, knockout, inhibitors, monoclonal antibodies, and nanobodies has significantly reduced angiogenesis, tumorigenesis, tumour growth, and metastasis and improved survival (7,23,28–37). Among inhibitors, EG00229 is the most specific NRP1 inhibitor, blocking its interaction with VEGFA and demonstrating tumor-suppressive effects in various cancers (28,38–42). While NRP1 is a potential target in melanoma, EG00229’s effects on melanoma remain unstudied (43,44). Additionally, although NRP1 depletion and inhibition have shown favourable anti-tumour immune responses in cancer cells and intra-tumoral Tregs, its impact on peripheral Tregs is less explored but promising (35–37,45). Also, the potential anti-tumour role of EG00229 has been studied in different cancers (41,46,47), but its effect on Tregs (intratumoral or peripheral) remains unexplored.

This study explores how Treg-mediated suppression of peripheral splenic T cells in B16-F10 melanoma is affected by NRP1 inhibition using EG00229 trifluoroacetate (EG). It uncovers the underlying cellular pathways, immune regulation, and anti-tumor responses. These findings suggest NRP1 as a critical checkpoint in peripheral Tregs and a potential target for cancer immunotherapy, alone or in combination with standard treatments.

## Materials and Method

### Cell lines

The murine melanoma cell line B16-F10 (ATCC™ CRL-6475™) and the murine fibroblast cell line NIH-3T3 (ATCC™ CRL-1658™) were cultured in Dulbecco’s Modified Eagle’s Medium (DMEM; Gibco, Grand Island, New York) supplemented with 10% fetal bovine serum (FBS; Gibco), 1X antibiotic solution (100X Liquid, HiMedia Laboratories Pvt. Ltd., Mumbai, India), and 1X L-glutamine (HiMedia). The cells were incubated at 37°C in a humidified CO_2_ incubator. For passaging, 1X trypsin solution (HiMedia) was used for both B16-F10 and NIH-3T3 cells.

### Antibodies and Reagents

PE Annexin V Apoptosis Detection Kit I, purified NA/LE Hamster Anti-mouse CD3e (cat no.-567114) and CD28 (cat no.-553294) antibodies, fluorophore-conjugated anti-mouse CD3 (cat no.- 553062), CD279 (PD-1, cat no.-J43), CD274 (PD-L1, cat no.-M1H5), NKG2AB6 (cat no.-16a11), CD152 (CTLA-4, cat no.-UC10-4F10-11), CD4 (cat no.-GK1.5) antibodies were from BD Biosciences (SJ, USA). Fluorophore-conjugated anti-mouse CD304 (NRP1, cat no.- 25-3041-82), FOXP3 (cat no.-FJK-16s), GATA-3 (cat no.-TWAJ) and T-bet (4B10) antibodies, Dynabeads™ Untouched™ Mouse T Cells Kit, CellTrace CFSE Cell Proliferation Kit, Foxp3 / Transcription Factor Staining Buffer Set, and SYBR Green qPCR Master Mix were from ThermoFisher Scientific (Waltham, USA). Anti-mouse Phospho-P38 MAPK (cat no.-4511s), P38 MAPK (cat no.-9212s), Phospho-P44/42 MAPK (Erk1/2) (cat no.-4370s), P44/42 MAPK (Erk1/2) (cat no.-4695BC), Phospho-SAPK/JNK (Thr183/Tyr185) (cat no.-4668s), SAPK/JNK (cat no.-9252s), Phospho-STAT3 (Tyr705) (cat no.-9145s), STAT3 (cat no.-9172s), Phospho-STAT1 (cat no.-8826s), STAT1 (cat no.- 9172s), Phospho-STAT5 (cat no.-9359s), STAT5 (cat no.-94205s), Phospho-Smad2/3 (cat no.-8828s), Smad2/3 (cat no.-8685s), Phospho-AKT (Thr308) (cat no.-13038T), AKT (cat no.-4691T), Phospho-PI3K (cat no.-17366), PI3K (cat no.-4257), and β-actin (cat no.-8457) antibodies and RIPA buffer were from Cell Signalling Technology (CST, Danvers, MA, USA). Saponin, Bradford reagent, NP-40, sodium deoxycholate, sodium dodecyl sulphate (SDS), protease inhibitor cocktail, and bovine serum albumin fraction V (BSA) were purchased from Sigma-Aldrich. PVDF membrane and Immobilon Western Chemiluminescent HRP substrate were purchased from Millipore (MA, USA). Sodium chloride and Tris base were purchased from HiMedia. TRIzol reagent was from Invitrogen (MA, USA). PrimeScript 1st strand cDNA Synthesis Kit was from Takara Bio (USA). Primers for Real-time PCR were from GCC Biotech India Pvt Ltd (West Bengal, India). EG00229 trifluoroacetate (an inhibitor for NRP1, cat no.- 6986) was from Tocris (United Kingdom).

### Cell culture supernatant collection

B16-F10 cells were cultured in DMEM following the ATCC protocol (48,49). When the cells reached 60–70% confluence, the culture medium was replaced with a fresh medium. After 24 hours (24h), the cell-free supernatant, or conditioned medium (B16-F10-CS), was collected and stored at −80°C until further use (50,51).

### Trypan blue exclusion test of cell viability

Purified murine splenic T cells were incubated for 96 hours with varying concentrations of EG with or without TCR activation. After incubation, the cells were harvested and washed with 1X PBS. Trypan blue was added, and the viable cells were manually counted, as described earlier (50). For the positive control, T cells were either heat-killed (65°C for 5 minutes in a water bath) or exposed to UV light (30 mins) before performing the cell viability assay.

### Mice

Female C57BL/6 mice, aged 6 to 8 weeks, were obtained from the National Institute of Science Education and Research (NISER), Bhubaneswar, for experimentation. The Institutional Animal Ethics Committee (IAEC) approved the study protocols at NISER, adhering to the guidelines established by the Committee for Control and Supervision of Experiments on Animals (CCSEA).

### B16-F10 cell injection (subcutaneously or intravenously), EG treatment, tumour volume measurement, metastatic lung nodule count and survivability assay

Five different groups, each group containing 5 mice, were made. The five groups made were: control, B16-F10 tumour-bearing, EG-treatment in B16-F10 tumour-bearing, B16-F10 intravenously injected and EG-treatment in B16-F10 intravenously injected. B16-F10 cells were injected subcutaneously or intravenously according to the earlier protocol (52). B16-F10 cells in their logarithmic growth phase were collected using trypsin when the flasks were no more than 50% full. The cells were mixed in ice-cold 1X PBS at 0.1 million cells/mL concentrations for subcutaneous injections and 0.2 million cells/mL for intravenous injections. For subcutaneous injection, 100 μL of the cell solution was injected into the upper right flank of the mice using an insulin syringe, ensuring a small bump formed. The control group received the same volume of chilled 1X PBS. For intravenous injections, mice had their tails dipped in warm water to enlarge the veins, and 0.5 mL of the cell solution was injected into their tail veins. The control group also received the same volume of chilled 1X PBS.

EG treatment in the B16-F10 tumour-bearing group was given as described earlier (53,54). Briefly, In the B16-F10 tumour-bearing group, EG treatment (10 mg/kg body weight, 100 μL per mouse) was given three times a week for two weeks, starting on the 14th day after subcutaneous injection, once the tumours became noticeable (about 10-12 days post-injection). Mice in control and B16-F10 tumour-bearing groups were injected with 1X PBS. Tumour growth was measured periodically from day 14 using a digital calliper (Fisher Scientific), and tumour volume was calculated with the formula [(tumour length × tumour width²)/2]. Afterwards, all mice were euthanised via CO[ asphyxiation, and their spleens, along with subcutaneous tumours from tumour-bearing mice (with or without EG treatment), were harvested for analysis. EG treatment in the B16-F10 intravenously injected group was given as described earlier (53,54). In the B16-F10 intravenously injected group, EG treatment (10 mg/kg, 100 μL/mouse) was administered intraperitoneally three times a week for three weeks, starting the second-day post-injection. Control and B16-F10 groups received 1X PBS. Afterwards, all mice were euthanised via CO[ asphyxiation, and the lungs and heart were removed as a single unit. The lungs were rinsed with water 1X PBS and metastatic lung nodules were counted on both lobes using a lamp and magnifying glass.

In a separate group of mice, including B16-F10 tumour-bearing, EG-treated B16-F10 tumour-bearing, B16-F10 intravenously injected, and EG-treated B16-F10 intravenously injected groups, the same injection and treatment regimens were followed without sacrificing the mice. Survival rates were assessed. For subcutaneous survival experiments, mice were killed when tumours reached 20 mm^3^ in size or tumour necrosis exceeded 50% of the tumour surface.

### T cell isolation, purification, and cell culture

Around 1.5 million purified murine splenic T cells were cultured in each experimental condition. First, splenocytes from mice were isolated as described elsewhere (55). In brief, spleens from (either control, B16-F10 tumour-bearing or B16-F10 tumour-bearing with EG treatment) C57BL/6 mice were disrupted through a 70 µM cell strainer. RBCs were lysed using RBC lysis buffer, washed with 1X PBS, centrifuged, and resuspended in complete RPMI-1640. T cells were then purified using the Untouched T cell Isolation Kit according to the manufacturer’s protocol (50). Splenocytes were resuspended in an isolation buffer and incubated with biotinylated antibodies. Cells were then washed with excess isolation buffer, centrifuged, and incubated for 15 minutes (mins) with streptavidin-conjugated magnetic beads and then placed in a magnet for 2 mins. The purity of T cells was≥ 95%, measured using flow cytometry (BD LSRFortessa™ Cell Analyzer) and analysed by the FlowJo™ v10 software. T cell activation assays were conducted using anti-CD3 (plate-bound) and CD28 antibodies (TCR activation) with or without B16-F10-CS (constituting 5% of the total volume of the cell suspension and media in the experimental setup) and EG00229 trifluoroacetate (EG) at a concentration of 7μM, as previously described. EG concentration of 7 μM in all *in vitro* experimental conditions was used as per earlier studies (41,56) as well as toxicity assay and subsequent estimation of EC_50_ value of EG on purified murine splenic T cells (Supplementary figure S1). The cells were incubated for 96h at 37°C in a humidified CO_2_ incubator. For western blot analysis, T cells were cultured with or without B16-F10-CS and EG for 96h and then subjected to TCR activation or left untreated for 15 mins before harvesting.

### Flow Cytometry (FC)

Flow cytometric assay was performed as mentioned elsewhere (57,58). Around 1.5 million purified murine splenic T cells were cultured in each experimental condition. In brief, purified T cells from different experimental conditions were suspended in FACS buffer (1X PBS, 1% BSA, 0.01% NaN3) were stained with fluorophore-conjugated antibodies against specific cell surface markers (CD3, CD4, NRP1, NKG2A) in unfixed condition, incubated for 30 min on ice in dark, and washed twice with ice-cold FACS buffer to remove unbound antibodies and suspended in 1% PFA and kept at 4[ for further analysis. Following the manufacturer’s protocol, intercellular staining for FOXP3 was done using Foxp3/Transcription Factor Staining Buffer Set. Briefly, purified T cells stained with fluorophore-conjugated anti-mouse CD3 and CD4 antibodies were washed with FACS buffer and incubated with 1X permeabilisation/fixation (perm/fix) buffer for 45mins at room temperature (RT) in the dark. Cells are then washed with 1X permeabilisation (perm) buffer, incubated with anti-mouse fluorophore-conjugated FOXP3 antibody, and incubated for 45 minutes at RT in the dark. Cells were then washed with perm buffer, suspended in 1% PFA and stored at 4[ for further analysis. Intercellular staining of CTLA-4, GATA-3 and T-bet was performed as described elsewhere (57,58). In brief, purified T cells stained with fluorophore-conjugated anti-mouse CD3 and CD4 antibodies were permeabilized with permeabilization buffer (1X PBS + 0.5% BSA + 0.1% Saponin + 0.01% NaN_3_) followed by blocking in 1% BSA (in permeabilization buffer) for 30mins at RT in dark. Then, the cells were washed with permeabilization buffer and incubated with anti-mouse fluorophore-conjugated antibodies against respective intercellular markers for 30 minutes at RT in the dark. Cells were then washed with permeabilization buffer, suspended in 1% PFA and stored at 4[ for further analysis. All the cells were acquired by the BD LSRFortessa™ Cell Analyzer (BD Biosciences) and analysed by the FlowJo™ v10 software (BD Biosciences). The gating strategy was as follows: the CD3^+^ positive population was selected from the total cell population shown in the FSC vs SSC dot plot. From the CD3^+^ positive population, the CD4^+^ cell population was gated (Supplementary figure S5). Within the CD3^+^ CD4^+^ dual positive population, the frequency of cells expressing various cellular markers or the expression levels of these markers were measured. The percentage of positive cells expressing the markers was referred to as frequency. In contrast, the expression levels of the markers on cells under different experimental conditions were represented by mean fluorescence intensity (MFI) values. The absolute count of CD4^+^ NRP1^+^ Tregs was calculated from the flow cytometric analysis using FlowJo software. Approximately ten thousand cells were acquired per sample.

### Apoptosis assay

Following the manufacturer’s protocol, flow cytometry (FC) was performed using the BD Annexin V Detection Kit I to detect apoptotic cells. Briefly, 1 million B16-F10 or NIH-3T3 cells were harvested and washed twice in ice-cold 1XPBS. Cells were then resuspended in 100 μL of 1X Annexin V binding buffer. PE-conjugated Annexin V and 7-AAD cocktail were added to each sample in the appropriate amounts, gently mixed by vortexing, and incubated at room temperature for 15 minutes in the dark. Afterwards, 400 µL of 1X Annexin V binding buffer was added to each tube. The samples were immediately acquired using the BD LSRFortessa™ Cell Analyzer and analysed with FlowJo™ v10 software.

### T cell Proliferation Assay

Purified T cells are incubated with 5 µM CFSE using CellTrace CFSE Cell Proliferation Kit following the manufacturer’s protocol, in 1X PBS for 10[minutes at room temperature and then washed three times with RPMI media supplemented with 10% FBS. Then, these CFSE-labelled T cells were cultured in different experimental conditions (Rest, TCR, B16-F10-CS, TCR+B16-F10-CS, TCR+EG and TCR+B16-F10-CS+EG) and incubated in a humified CO_2_ incubator for 96h. Also, purified splenic T cells isolated from either control, B16-F10-tumour-bearing or EG-treated B16-F10 tumour-bearing mice were stained with CFSE, washed times with RPMI media supplemented with 10% FBS and incubated in a humified CO_2_ incubator for 96h. Around 1.5 million purified murine splenic T cells were cultured in each experimental condition. Post incubation, cells from all experimental conditions were harvested, washed with 1XPBS and stained with fluorophore-conjugated anti-mouse CD3 and CD4 antibodies for 30 mins in the dark at 4[. Cells were then washed with FACS buffer, suspended in 1% PFA and stored at 4[ for further analysis. All the cells were acquired by the BD LSRFortessa™ Cell Analyzer (BD Biosciences) and analysed by the FlowJo™ v10 software (BD Biosciences). The gating strategy was as follows: the CD3^+^ positive population was selected from the total cell population shown in the FSC vs SSC dot plot. From the CD3^+^ positive population, the CD4^+^ cell population was gated. Within the CD3^+^ CD4^+^ dual positive population, the MFI of CFSE-stained cells was calculated.

### *In vitro* suppression assay of Tregs

The immunosuppressive potential of Tregs was evaluated through an *in vitro* suppression assay (37). Purified splenic T cells from control, B16-F10 tumor-bearing, and B16-F10 tumor-bearing mice treated with EG were cultured for 96 hours in a CO[ incubator, either with or without TCR stimulation, B16-F10-CS, or EG. Each experimental condition involved about 1.5 million T cells. Forty-eight hours before the end of this incubation, approximately 10 million T cells from normal C57BL/6 mice were activated using anti-CD3 and CD28 antibodies for 36-48 hours, then stained with CFSE. These activated T cells were co-cultured with the previously prepared T cells under various conditions (Rest, TCR, B16-F10-CS, TCR+B16-F10-CS, TCR+EG, and TCR+B16-F10-CS+EG) in a 1:1 ratio at a total density of 1.5 million cells/mL for 96 hours. After incubation, all cells were harvested, washed with PBS, and stained with fluorophore-conjugated anti-mouse CD3 and CD4 antibodies for 30 minutes in the dark at 4°C. The cells were then washed, suspended in 1% paraformaldehyde, and stored at 4°C for further analysis.

Flow cytometry was performed using a BD LSRFortessa™ Cell Analyzer, and data was analysed with FlowJo™ v10 software. The gating strategy involved selecting the CD3^+^ population from the total cell population and then identifying the CD4^+^ population within the CD3^+^ cells. The mean fluorescence intensity (MFI) of the CFSE-stained cells in the CD3^+^ CD4^+^ population was calculated to determine the proliferation rate of CFSE-labelled TCR-activated effector T cells in the presence of cells from control, experimental, or tumour-bearing mice with or without EG treatment.

### BD Cytometric Bead Array

To measure cytokine levels in purified T cells from control, B16-F10 tumor-bearing, and B16-F10 tumor-bearing mice treated with EG, the cells were cultured for 96 hours in a CO[ incubator with or without TCR stimulation, B16-F10-CS, or EG. Approximately 1.5 million T cells were used for each condition. The BD Cytometric Bead Array (CBA) Mouse Th1/Th2/Th17 Cytokine Kit was employed according to the manufacturer’s instructions (59). First, lyophilised standards were reconstituted with assay diluent and serially diluted in labelled tubes to create a standard curve, with a negative control containing only assay diluent. The Capture Beads were mixed, and aliquots were added to the assay tubes along with the cytokine standards and samples. After adding the detection reagent, the tubes were incubated for 2 hours at room temperature in the dark. Following incubation, wash buffer was added, and the tubes were centrifuged to discard the supernatant. Finally, the beads were resuspended in wash buffer and analysed immediately using the BD LSRFortessa™ Cell Analyzer and FlowJo™ v10 software per the manufacturer’s guideline (59).

### Western Blot Analysis

Protein expression was assessed by western blot analysis following previously described protocols (57). Purified T cells from control, B16-F10 tumor-bearing, and B16-F10 tumor-bearing mice treated with EG were cultured for 96 hours in a humidified CO[ incubator with or without TCR stimulation, B16-F10-CS, or EG. Approximately 10 million T cells were used for each condition. After incubation, the cells were washed with ice-cold PBS, and whole cell lysate (WCL) was prepared using RIPA lysis buffer and centrifuged at 13,000 rcf for 30 minutes at 4 °C. Protein concentrations were measured using the Bradford assay. Equal protein amounts were mixed with 6X Laemmli buffer, loaded onto 10% SDS-PAGE gels, and transferred to PVDF membranes. Membranes were blocked with 3% BSA in TBST and incubated overnight with anti-mouse primary antibodies (1:1000 dilution) targeting various proteins, including phosphorylated and total forms of MAPK, STAT, Smad, AKT, and PI3K, along with β-actin. After washing, HRP-conjugated secondary antibodies (1:4000 dilution) were applied for 2 hours at room temperature. The membranes were washed again and visualised using chemiluminescence detection with Immobilon Western HRP substrate. Images were captured using the Bio-Rad Gel Doc and analysed with Image Lab software, normalising against the β-actin as loading control.

### 2.13 cBioPortal analysis

The potential alterations of the NRP1 gene in previously reported cancer patient databases were analysed using the online server cBioPortal (https://www.cbioportal.org/, accessed on 6 June 2024). cBioPortal analysis was done in 212 melanoma cancer patients from a combined analysis of three studies: Metastatic Melanoma (DFCI, Science 2015), Melanoma (MSK, NEJM 2014), and Metastatic Melanoma (UCLA, Cell 2016) (Figure 2A). Additionally, 2683 cancer patient samples from a Pan-cancer study of whole genomes (ICGC/TCGA, Nature 2020) were also evaluated in the cBioPortal.

### Statistical analysis

Statistical analysis was conducted using GraphPad Prism 8.0 (GraphPad Software Inc., San Diego, CA, USA). Data are presented as Mean ± SEM. Unless specified otherwise, comparisons between groups were performed using One-way ANOVA or Student’s T-test. Using GraphPad Prism, the EC50 value of EG on purified murine splenic T cells was calculated from a non-linear dose-response curve. One-column T-test and Wilcoxon test were employed to assess the statistical significance of fold changes in gene expression between tumorigenic and non-tumorigenic cells. The significance of Kaplan–Meier survival curves for C57BL/6 mice injected with B16-F10 cells, either subcutaneously or intravenously with or without EG treatment, was evaluated using the Log Rank test in GraphPad Prism. All data are representative of at least three independent experiments, with P < 0.05 indicating statistical significance.

## Results

### Elevation of regulatory T cells (Tregs) frequency in murine splenic population during B16-F10 cell culture supernatant (B16-F10-CS) treatment, *in vitro*

The predominance of intratumoral Tregs and their role in immunosuppression, cancer progression, metastasis, and poor patient survival outcomes has been extensively reported (9,10,60). It has also been noted that cancer cell-conditioned media (CM) induces the generation of Tregs *in vitro*, facilitating cancer-mediated immunosuppression (50,61–63). Furthermore, while numerous studies show elevated expression of immunosuppressive markers in intratumoral Tregs, research on the expression and regulatory role of peripheral Tregs in cancerous environments or presence is limited (37,64,65). Accordingly, the current study assessed the possible increase in peripheral Tregs in the murine splenic T cell population in vitro during B16-F10-CS treatment (**Figure 1**).

**Figure 1:**
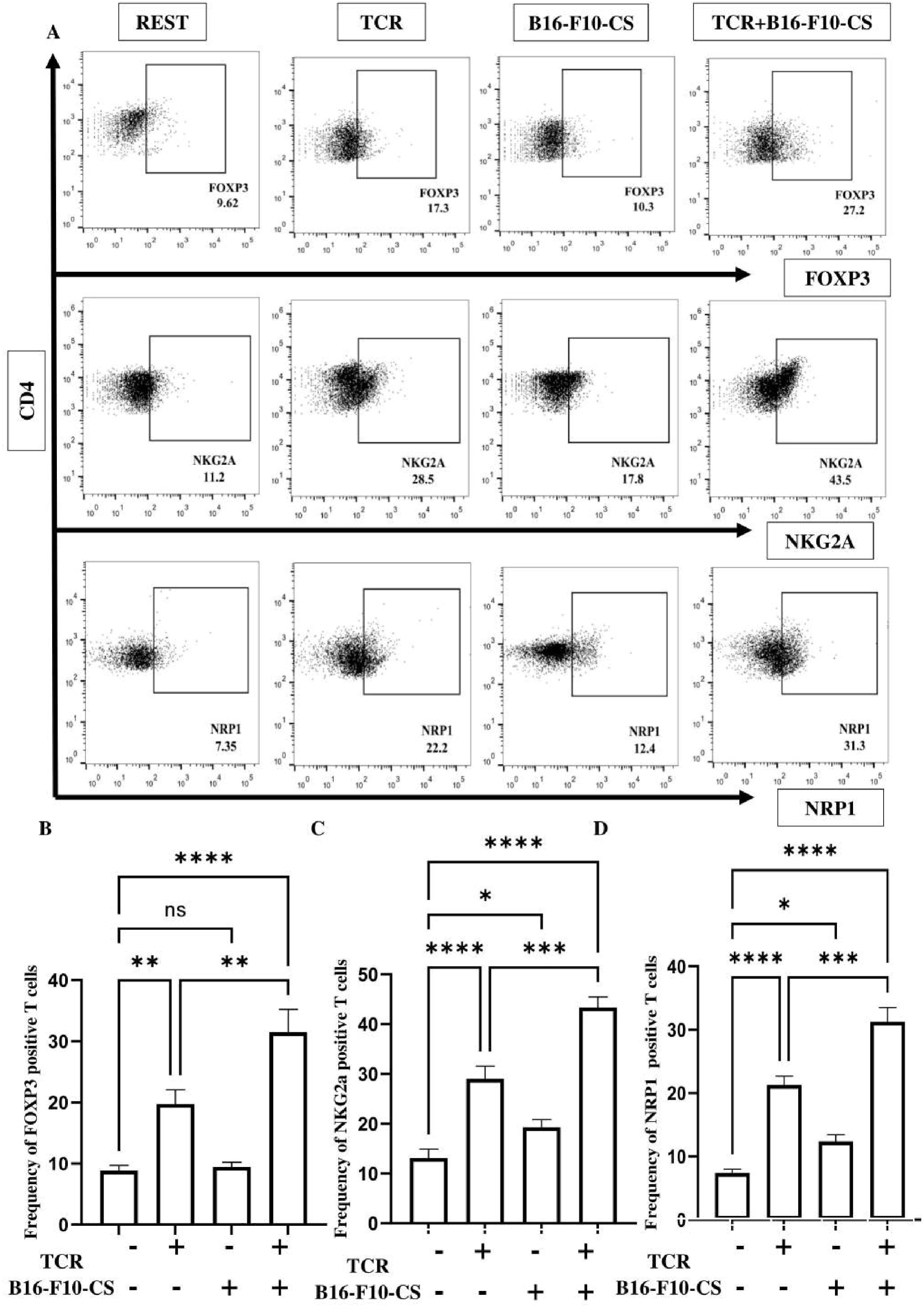
Elevation of frequency of suppressor T cells during B16-F10 cell culture supernatant (B16-F10-CS) treatment, *in vitro*. (A) Flow cytometric dot plots showing alterations in the frequency of FOXP3, NKG2A and NRP1 positive purified splenic T cell population with or without anti-CD3/CD28 antibodies and B16-F10-CS treatment. Bar graph representation of alteration in frequency of (B) FOXP3, (C) NKG2A and (D) NRP1 positive purified splenic T cell population with or without anti-CD3/CD28 antibodies and B16-F10-CS treatment. Representative bar diagrams are of three independent experiments. ns, non-significant; * p < 0.05; ** p < 0.01; *** p < 0.001; **** p < 0.0001.

**Figure 2:**
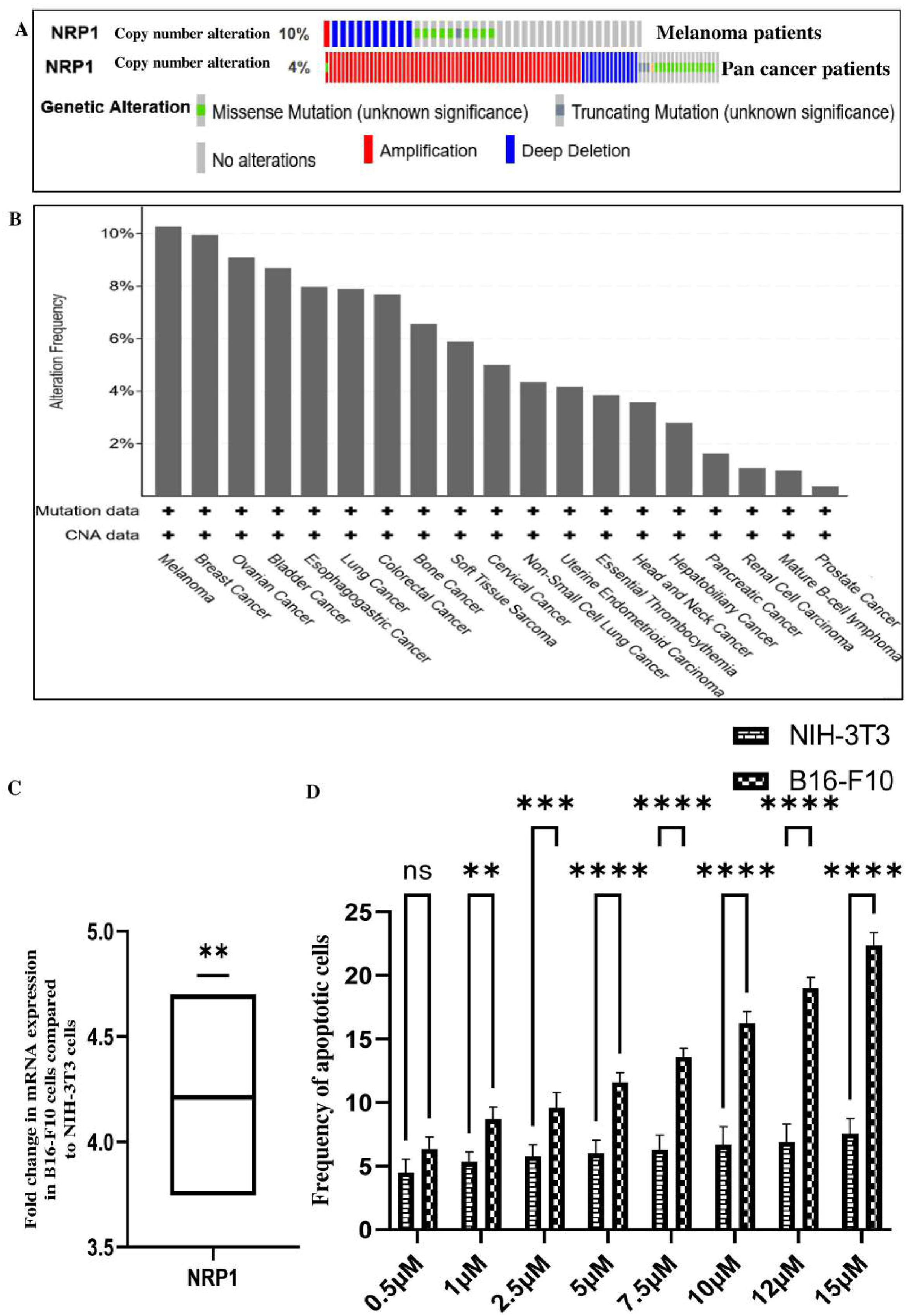
Neuropillin-1 (NRP1) expression is upregulated in melanoma patients and cell lines, and its inhibition induced apoptosis in tumorigenic B16-F10 cells. (A) Oncoprint depicting the frequency and types of alterations of the NRP1 gene in 212 melanoma cancer and 2683 pan-cancer patient samples from the cBioPortal database. (B) Bar graph representation of alteration frequency of NRP1 gene in 2683 cancer patient samples from cBioPortal database. (C) Fold change in mRNA expression of NRP1 gene when compared between tumorigenic B16-F10 and non-tumorigenic NIH-3T3 cells. (D) Bar graphs representing apoptosis in tumorigenic B16-F10 and non-tumorigenic NIH-3T3 cells upon treatment with different concentrations of NRP1 inhibitor EG00229, which is determined by Annexin & 7-AAD dual staining. Representative bar diagrams are of three independent experiments. ns, non-significant; * p < 0.05; ** p < 0.01; *** p < 0.001; **** p < 0.0001.

Flow cytometry dot plots showed that the frequency of FOXP3, NKG2A, and NRP1 positive T cells increased in the TCR+B16-F10-CS treatment compared to resting, TCR-activated, and B16-F10-CS-treated groups (**Figure 1A**), indicating an increase in the suppressor T cell population in the immunosuppressive B16-F10-CS condition. No significant change in the frequency of FOXP3-expressing T cells was found between resting and B16-F10-CS-treated T cells (**Figure 1B**), but the frequency of NKG2A and NRP1 positive T cells was higher in the B16-F10-CS-treated group compared to resting T cells (**Figures 1C and D**). While the frequency of other Treg markers like CTLA-4, PD-1, and PD-L1 was assessed, no noticeable differences were observed across the experimental conditions with TCR and TCR+B16-F10-CS treatments (**Supplementary figure S2**). Overall, the frequency of FOXP3, NKG2A, and NRP1 positive suppressor T cells increased in TCR-activated T cells after B16-F10-CS treatment *in vitro*.

### cBioPortal and mRNA analysis revealed that melanoma patients and cell lines had higher Neuropillin-1 (NRP1) expression and inhibiting NRP1-induced apoptosis in tumorigenic B16-F10 cells

Our previous data showed that the frequency of FOXP3, NKG2A, and NRP1 positive suppressor T cells increased during immunosuppressive B16-F10-CS treatment (**Figure 1**). NRP1 is a marker for the stability and functionality of regulatory suppressor T cells, and its depletion can reduce Treg suppressive activity (26). It has also been reported that carcinoma cell lines, including melanoma, rely on NRP1 for survival and growth by activating multiple growth factor receptors (44,47,66). Additionally, NRP1 expression in tumour-associated blood vessels and various cancers is linked to tumor progression, aggressiveness, advanced stages, and poor prognosis, particularly in melanoma and breast cancer (23,47,67). Therefore, the expression of NRP1 in cancer patients, including those with melanoma, was analysed using cBioPortal, while qRT-PCR was used to quantify NRP1 mRNA in tumorigenic B16-F10 and non-tumorigenic NIH-3T3 cell lines. We also evaluated the potential apoptotic effects of inhibiting NRP1 in these cell lines (**Figure 2**).

The potential changes in the NRP1 gene were analysed using the online server cBioPortal (https://www.cbioportal.org/, accessed on 6 June 2024). The oncoprint from cBioPortal showed that copy number alteration (CNA) of the NRP1 gene was found in 10% of 212 melanoma patients based on a combined analysis of three studies: Metastatic Melanoma (DFCI, Science 2015), Melanoma (MSK, NEJM 2014), and Metastatic Melanoma (UCLA, Cell 2016) (**Figure 2A**). Additionally, the oncoprint indicated that CNA of the NRP1 gene was found in 4% of 2,683 cancer patient samples from a Pan-cancer study of whole genomes (ICGC/TCGA, Nature 2020). Among various mutation types, NRP1 gene amplification was observed in melanoma and other cancer patients. The bar graph showed that the frequency of NRP1 gene alterations was highest in melanoma patients based on the CNA and mutation load data from the Pan-cancer study (**Figure 2B**).

To confirm the cBioPortal findings of increased NRP1 expression in melanoma and other cancer patients, mRNA levels of NRP1 were compared between tumorigenic B16-F10 and non-tumorigenic NIH-3T3 cells. NRP1 expression was higher in B16-F10 cells than in NIH-3T3 cells (**Figure 2C**).

Since NRP1 is significantly involved in melanoma cell viability and proliferation, we evaluated the effect of NRP1 inhibition in B16-F10 and NIH-3T3 cells using the specific small molecule inhibitor EG00229 trifluoroacetate (EG). We assessed apoptosis with Annexin V and 7-AAD dual staining. Flow cytometry dot plots and representative bar graphs showed minimal necrotic cell death (indicated by 7-AAD positive cells) in both B16-F10 and NIH-3T3 cells treated with different concentrations of EG (**Figure 2D, Supplementary Figure S3**). However, there was a gradual increase in apoptotic B16-F10 cells (both early and late) with EG treatment at doses higher than 10 μM, while negligible cell death occurred in NIH-3T3 cells treated with EG. Thus, cBioPortal analysis revealed elevated NRP1 expression in cancer patients, including melanoma patients. The mRNA quantification showed heightened NRP1 levels in B16-F10 melanoma cells compared to the control cell line. Additionally, inhibiting NRP1 led to apoptosis in tumorigenic B16-F10 cells but not in non-tumorigenic NIH-3T3 cells.

### Inhibition of Neuropillin-1 (NRP1) decreased the expression of regulatory T cell (Treg) markers in splenic T cells treated with B16-F10 cell culture supernatant (B16-F10-CS) *in vitro*

In our previous results, we have observed elevated frequency of FOXP3, NKG2A and NRP1 positive Tregs in TCR-activated splenic T cells cultured in B16-F10-CS (**Figure 1**). Earlier studies suggest that NRP1-deficient Tregs retain FOXP3 expression, but depletion of NRP1 in T cells significantly decreases the frequency of tumour-infiltrating FOXP3^+^ Tregs in the TME (65,68,69). Also, NRP1 expression is observed in newly emigrated thymic natural killer T (NKT) cells, yet little is known about its role in these cells (70). Therefore, we wanted to study whether NRP1 plays any possible role in the modulation of other Tregs, such as FOXP3, NKG2A, CTLA-4, PD-1, and PD-L1 (64,71–75) during B16-F10-CS driven immunosuppressive conditions. Accordingly, we evaluated the expression of Treg markers (NRP1, FOXP3, and NKG2A, CTLA-4, PD-1, PD-L1) on TCR-activated B16-F10-CS-treated CD4^+^ splenic T cells, with or without NRP1 inhibition using EG (**Figure 3 and Supplementary Figure S4**).

**Figure 3:**
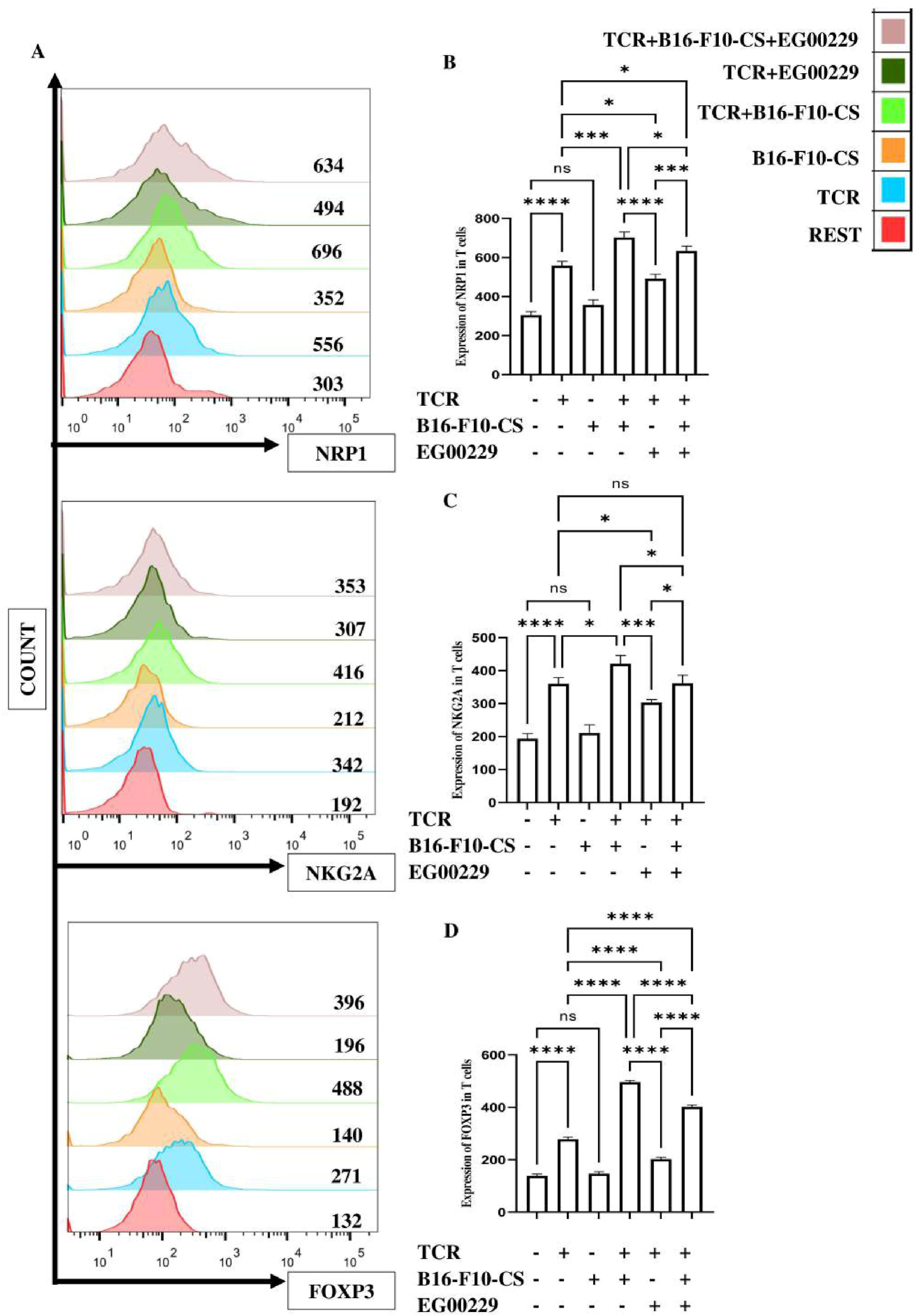
Inhibition of Neuropillin-1 (NRP1) alleviated expression of suppressor T cell markers in splenic T cells treated with B16-F10 cell culture supernatant (B16-F10-CS), *in vitro*. (A) Flowcytometric histogram plots depicting alterations in the expression of NRP1, NKG2A and FOXP3 in anti-CD3/CD28 antibodies and B16-F10-CS treated purified splenic T cell with or without NRP1 inhibition. Bar graph representation of alteration in expression of (B) NRP1, (C) NKG2A and (D) FOXP3 in anti-CD3/CD28 antibodies and B16-F10-CS treated purified splenic T cell with or without NRP1 inhibition. Representative bar diagrams are of three independent experiments. ns, non-significant; * p < 0.05; ** p < 0.01; *** p < 0.001; **** p < 0.0001.

The expression of immunosuppressive Treg markers (NRP1, NKG2A, and FOXP3) was increased in splenic T cells cultured under TCR+B16-F10-CS conditions. Treatment with EG reduced the expression of these Treg markers (**Figure 3A-D**). Similarly, elevated expression of these markers in TCR-activated splenic T cells decreased following EG treatment. Although the expression of other Treg markers, such as CTLA-4, PD-1, and PD-L1, was examined, no significant changes were observed between the TCR and TCR+B16-F10-CS treatment conditions (**Supplementary Figure S4**). Therefore, NRP1 inhibition with EG in CD4^+^ splenic T cells cultured in the immunosuppressive B16-F10-CS environment not only decreased NRP1 expression but also lowered NKG2A and FOXP3 levels *in vitro*.

### Neuropilin-1 (NRP1) inhibition increased proinflammatory cytokine secretion and T cell proliferation while reducing Treg suppressive activity and immunosuppressive cytokines release in peripheral Tregs during B16-F10 cell culture supernatant (B16-F10-CS) treatment *in vitro*

It has been reported earlier that NRP1 acts as a functionality and stability marker for Treg, and its depletion diminished Treg immunosuppressive potential, promoted Treg fragility and ameliorated anti-tumour response (7,23,31,76–78). Our previous data showed that NRP1 inhibition downregulated the expression of NRP1 and other Treg markers in B16-F10-CS-treated splenic T cells (**Figure 3**), indicating its probable role in immune modulation. Accordingly, we evaluated TCR-activated B16-F10-CS-treated CD4+ splenic T cells’ immunosuppressive potential with or without NRP1 inhibition with EG (**Figure 4**).

**Figure 4:**
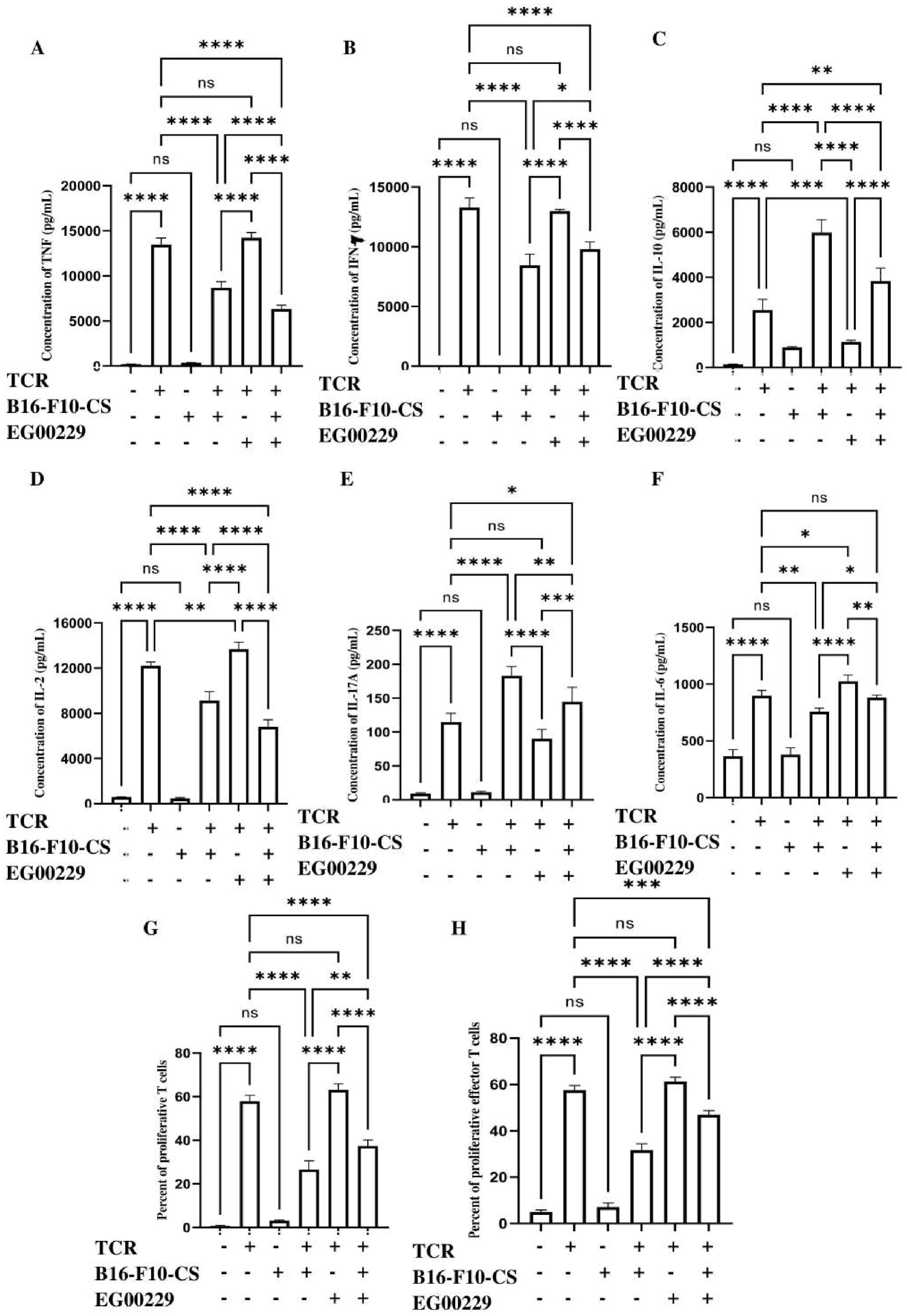
Inhibition of Neuropilin-1 (NRP1) enhanced proinflammatory cytokine secretion and T cell proliferation while decreasing Treg suppressive activity and the release of immunosuppressive cytokines in splenic T cells treated with B16-F10 cell culture supernatant (B16-F10-CS), *in vitro*. Bar graphs depicting the alteration in (A) TNF (B) IFN-γ, (C) IL-10, (D) IL-2, (E) IL-17(A), and (F) IL-6 secretion levels in anti-CD3/CD28 antibodies and B16-F10-CS treated purified splenic T cell with or without NRP1 inhibition. (G) Bar graph depicting proliferation of T cells via CFSE proliferation assay upon treatment with anti-CD3/CD28 antibodies and B16-F10-CS with or without NRP1 inhibition. (H) Bar graph depicting proliferation of effector T cells co-cultured with purified splenic T cells treated with anti-CD3/CD28 antibodies and B16-F10-CS with or without NRP1 inhibition. Representative bar diagrams are of three independent experiments. ns, non-significant; * p < 0.05; ** p < 0.01; *** p < 0.001; **** p < 0.0001.

CBA analysis for quantification of cytokines secreted from TCR+B16-F10-CS-treated splenic T cells revealed that effector cytokines (TNF, IFN-γ, IL-2, IL-6) (**Figure 4A, B, D and F**) release was attenuated. Alternatively, these cells showed augmented secretion of immunomodulatory cytokines (IL-10, IL-17A) (Figure 4C and E). This was reversed in TCR+B16-F10-CS+EG-treated splenic T cells. There was no significant change in TNF, IFN-γ, and IL-17A levels in only TCR-activated T cells with or without EG treatment. However, levels of IL-2 and IL-6 were increased, while IL-10 levels were decreased in TCR+EG-treated T cells compared to untreated TCR-activated ones.

Furthermore, the flow cytometric histogram from the CFSE assay and representative bar diagram depicted that cell proliferation was diminished in TCR+B16-F10-CS-treated splenic T cells. This reduced proliferative rate was ameliorated in TCR+B16-F10-CS+EG-treated splenic T cells (**Figure 4G and Supplementary figure S7A**). There was no discernible change in cell proliferation of TCR-activated T cells with or without EG treatment. Furthermore, the absolute CD4^+^ NRP1^+^ Treg count among 1.5 million purified murine splenic T cells cultured in different experimental conditions was evaluated (**Supplementary figure S6A and B**). The absolute count of CD4^+^ NRP1^+^ Tregs was increased in the TCR+B16-F10-CS condition compared to only the TCR-activated condition. NRP1 inhibition with EG showed diminished CD4^+^ NRP1^+^ Treg number compared between TCR and TCR+EG or TCR+B16-F10-CS and TCR+B16-F10-CS+EG conditions.

We further evaluated the immunosuppressive potential of CD4^+^ TCR+B16-F10-CS-treated splenic T cells with or without EG treatment. To address this, we co-cultured CFSE-labelled TCR-activated effector T cells with splenic T cells cultured in different experimental conditions (Rest, TCR, B16-F10-CS, TCR+B16-F10-CS, TCR+EG, TCR+B16-F10-CS+EG). Then, we assessed the cell proliferation of these effector T cells to evaluate the effect of the splenic T cells in different experimental conditions mentioned above. It was observed that TCR+B16-F10-CS+EG treated CD4^+^T cells increased the proliferation of co-cultured effector T cells compared to TCR+B16-F10-CS-treated T cells (**Figure 4H and Supplementary figure S7B**). No visible change was observed in the proliferation of effector T cells co-cultured with TCR-activated T cells with or without EG treatment.

Thus, it was observed that NRP1 inhibition with EG not only upregulated CD4^+^ T cell proliferation and effector cytokine response in TCR-activated B16-F10-CS-treated T cells but also downregulated secretion of immunomodulatory cytokines and Treg suppressive activity.

### Neuropilin-1 (NRP1) inhibition suppressed STAT, Smad2/3, and ERK-MAPK pathways while activating the AKT/PI3K pathway in peripheral Tregs during B16-F10 cell culture supernatant treatment *in vitro*

NRP1 in cancer cells and intratumoral Tregs is reported to regulate STAT, PI3K/AKT, TGF-β-mediated Smad2/3, and ERK MAPK signalling pathways during various cancers, thereby facilitating cancer progression and metastasis (34,75,79–83). Accordingly, we evaluated the effect of NRP1 inhibition with EG on STAT, PI3K/AKT, Smad2/3, and ERK-MAPK signalling in murine splenic peripheral Tregs cultured in immunosuppressive B16-F10-CS (**Figure 5**).

**Figure 5:**
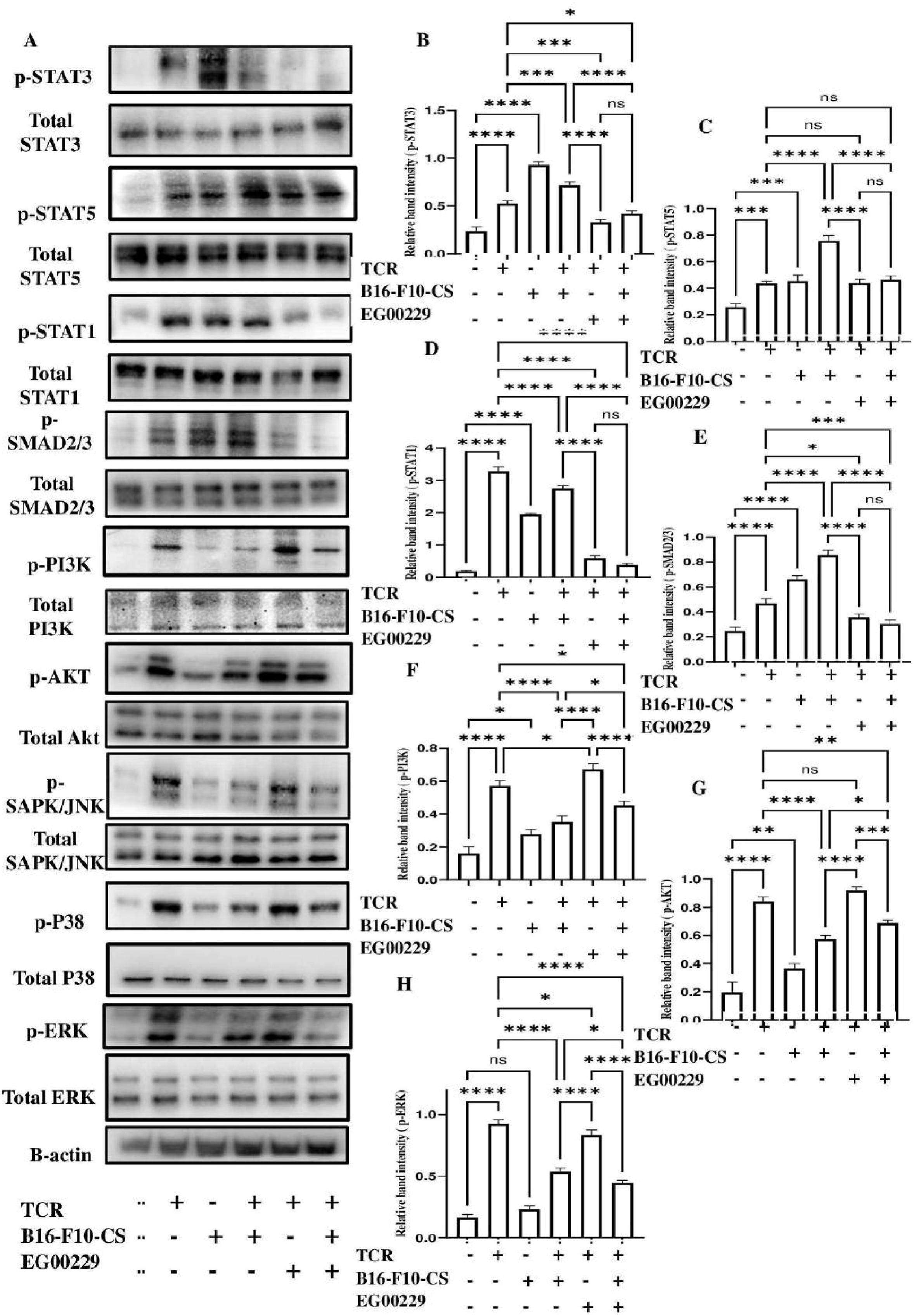
Inhibition of Neuropilin-1 (NRP1) alleviated the STAT, SMAD2/3, and ERK MAPK pathways while enhancing the AKT/PI3K pathway in splenic T cells treated with B16-F10 cell culture supernatant (B16-F10-CS), *in vitro*. (A) Western blots showing expression of phosphorylated STAT3, STAT5, STAT1, SMAD2/3, PI3K, AKT, SAPK/JNK, P38 and ERK in anti-CD3/CD28 antibodies and B16-F10-CS treated purified splenic T cell with or without NRP1 inhibition. Representative bar graphs are of three independent experiments, showing relative expression levels of (B) p-STAT3, (C) p-STAT5, (D) p-STAT1, (E) p-SMAD2/3, (F) p-PI3K, (G) p-AKT and (H) p-ERK when compared with expression of total STAT3, STAT5, STAT1, SMAD2/3, PI3K, AKT and ERK respectively. Β-actin was used as a loading control for this experiment. ns, non-significant; * p < 0.05; ** p < 0.01; *** p < 0.001; **** p < 0.0001.

Western blot analysis and representative bar graphs revealed that TCR+B16-F10-CS+EG-treated splenic T cells exhibited a diminished expression of phosphorylated STAT1, STAT3, STAT5, Smad2/3, and ERK as compared to TCR+B16-F10-CS-treated ones (**Figure 5A, B, C, D, E and H**). Conversely, TCR+B16-F10-CS+EG-treated T cells showed increased phosphorylated AKT and PI3K expression compared to only TCR+B16-F10-CS-treated ones (**Figure 5F and G**). No discernible alteration in phosphorylated STAT5 and AKT expression was observed in TCR-activated murine splenic T cells with or without NRP1 inhibition. However, the expression of phosphorylated STAT1, STAT3, Smad2/3, and ERK was reduced in TCR+EG-treated splenic T cells compared to only TCR-activated ones. On the contrary, PI3K phosphorylation was upregulated in TCR+EG-treated splenic T cells compared to only TCR-activated ones. No visible alteration in the expression of phosphorylated SAPK/JNK and P38 (**Figure 5A and Supplementary Figure S8**) was observed when compared between TCR-activated and TCR+EG-treated T cells or TCR+B16-F10-CS-treated and TCR+B16-F10-CS+EG treated T cells. This indicated that NRP1 might not directly regulate SAPK/JNK and P38 MAPK pathways in splenic peripheral Tregs cultured in B16-F10-CS *in vitro*.

Therefore, NRP1 inhibition with EG was observed to decrease STAT, Smad2/3 and ERK MAPK phosphorylation while enhancing AKT and PI3K phosphorylation in splenic peripheral Tregs treated with B16-F10-CS.

### Treatment with a Neuropilin-1 (NRP1) inhibitor decreased Treg and Th2 markers and increased Th1 marker expression in splenic T cells from B16-F10 tumor-bearing mice

It has been reported that intratumoral Tregs expressing NRP1, NKG2A, FOXP3, PD-1, PD-L1, CTLA-4, GATA-3, and T-bet play a vital role in the tumour environment, influencing tumour survival, growth, spread, and patient outcomes, making them targets for cancer immunotherapy (64,71–75). On the other hand, few studies explore the expression of these Treg markers in peripheral Tregs and their possible role in tumour growth and metastasis (76,84). While NRP1 inhibition or depletion is known to affect FOXP3, T-bet, CTLA-4, and PD-1 in intratumoral Tregs (23), its possible role in regulating these molecules in peripheral Tregs during cancer is still unknown. Accordingly, we evaluated the expression of Treg markers such as NRP1, NKG2A, FOXP3, PD-1, PD-L1, CTLA-4, GATA-3 and T-bet in murine splenic T cells isolated from B16-F10 melanoma tumour-bearing mice with or without NRP1 inhibitor (EG) treatment.

It was observed that splenic T cells isolated from EG-treated B16-F10 melanoma tumour-bearing mice showed attenuated expression of Treg markers such as NRP1, NKG2A, FOXP3, PD-1, PD-L1 and CTLA-4, and Th2 marker GATA-3 when compared to T cells isolated from untreated tumour bearing mice (**Figure 6A-G and Supplementary figure S9A-G**). Conversely, T-bet (Th1 marker) expression was elevated in splenic T cells isolated from EG-treated tumour-bearing mice compared to T cells isolated from untreated tumour-bearing mice (**Figure 6H and Supplementary figure S9H**).

**Figure 6:**
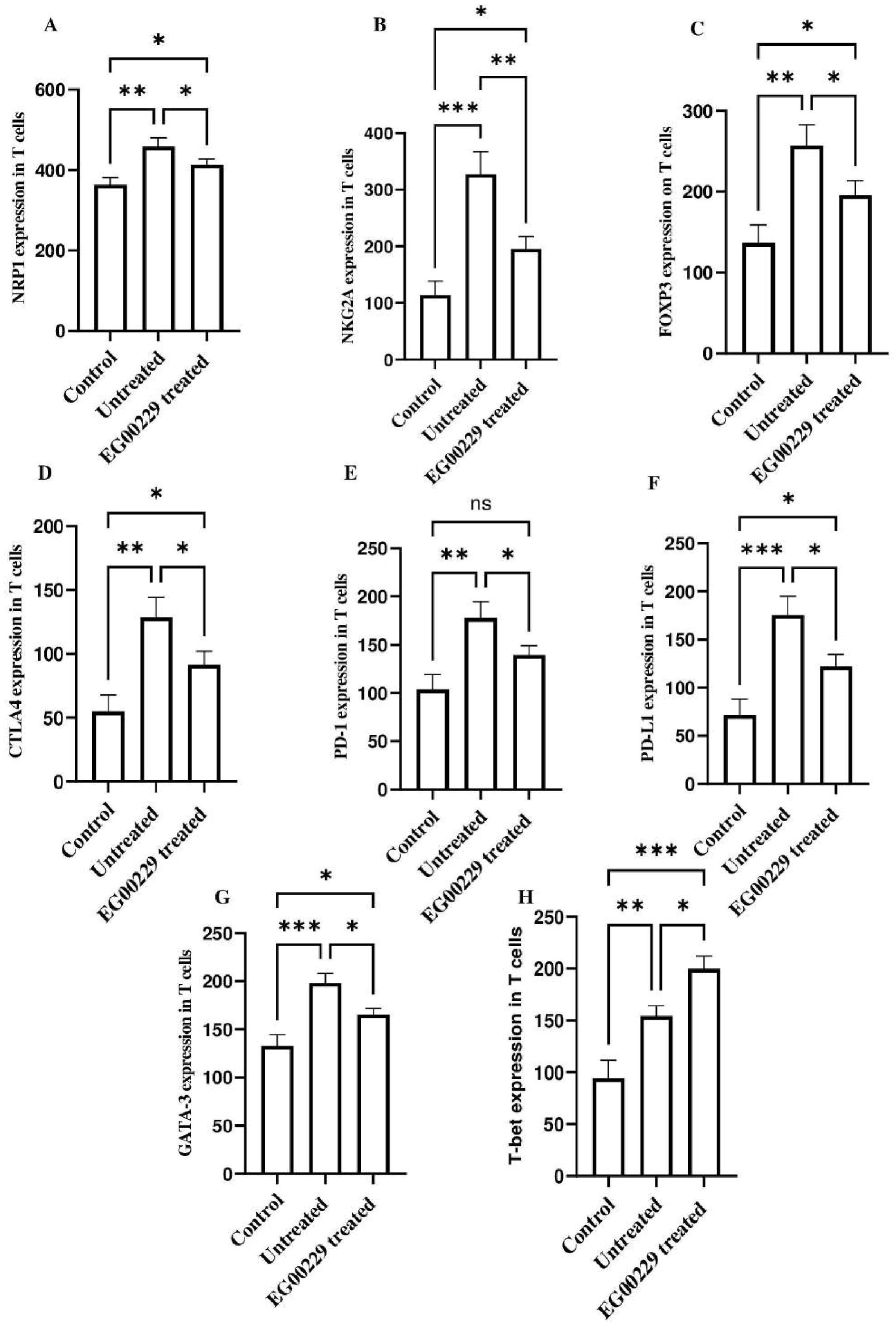
Expression of suppressor T cell markers alleviated in splenic T cells isolated from B16-F10 subcutaneous tumour-bearing mice treated with Neuropillin-1 (NRP1) inhibitor. Bar graphs depicting the alteration in expression of (A) NRP1, (B) NKG2A, (C) FOXP3, (D) T-bet, (E) GATA-3, (F) CTLA-4, (G) PD-1 and (H) PD-L1 in purified splenic T cells isolated from control healthy and B16-F10 subcutaneous tumour bearing C57BL/6 mice with or without NRP1 inhibitor treatment. Representative bar diagrams are of three independent experiments. ns, non-significant; * p < 0.05; ** p < 0.01; *** p < 0.001; **** p < 0.0001.

Thus, it was observed that peripheral splenic T cells from EG-treated subcutaneous B16-F10 melanoma tumour-bearing mice showed decreased expression of Treg markers such as NRP1, NKG2A, FOXP3, CTLA-4, PD-1, PD-L1, and the Th2 transcription factor GATA-3. These T cells also showed elevated expression of Th1 transcription factor T-bet.

### Neuropilin-1 (NRP1) inhibitor treatment increased proliferation and reduced immunosuppressive activity of splenic T cells from B16-F10 tumor-bearing mice

NRP1 is reported to negatively regulate the proliferation of CD4^+^FOXP3^+^ Tregs (7). It has also been reported that NRP1 inhibits the suppressive function of intratumoral regulatory T cells (37). Furthermore, NRP1 expression on FOXP3^+^ Tregs is crucial for T cell suppression by regulating conventional T cell proliferation, phenotype, cytokine production, and suppressive function *in vitro* and *in vivo* (85). Accordingly, the proliferation of CD4^+^ splenic T cells isolated from B16-F10 subcutaneous tumour-bearing mice, with or without EG treatment, and their suppressive activity on the TCR-activated splenic effector T cells were evaluated by the CFSE cell proliferation assay (**Figure 7**).

**Figure 7:**
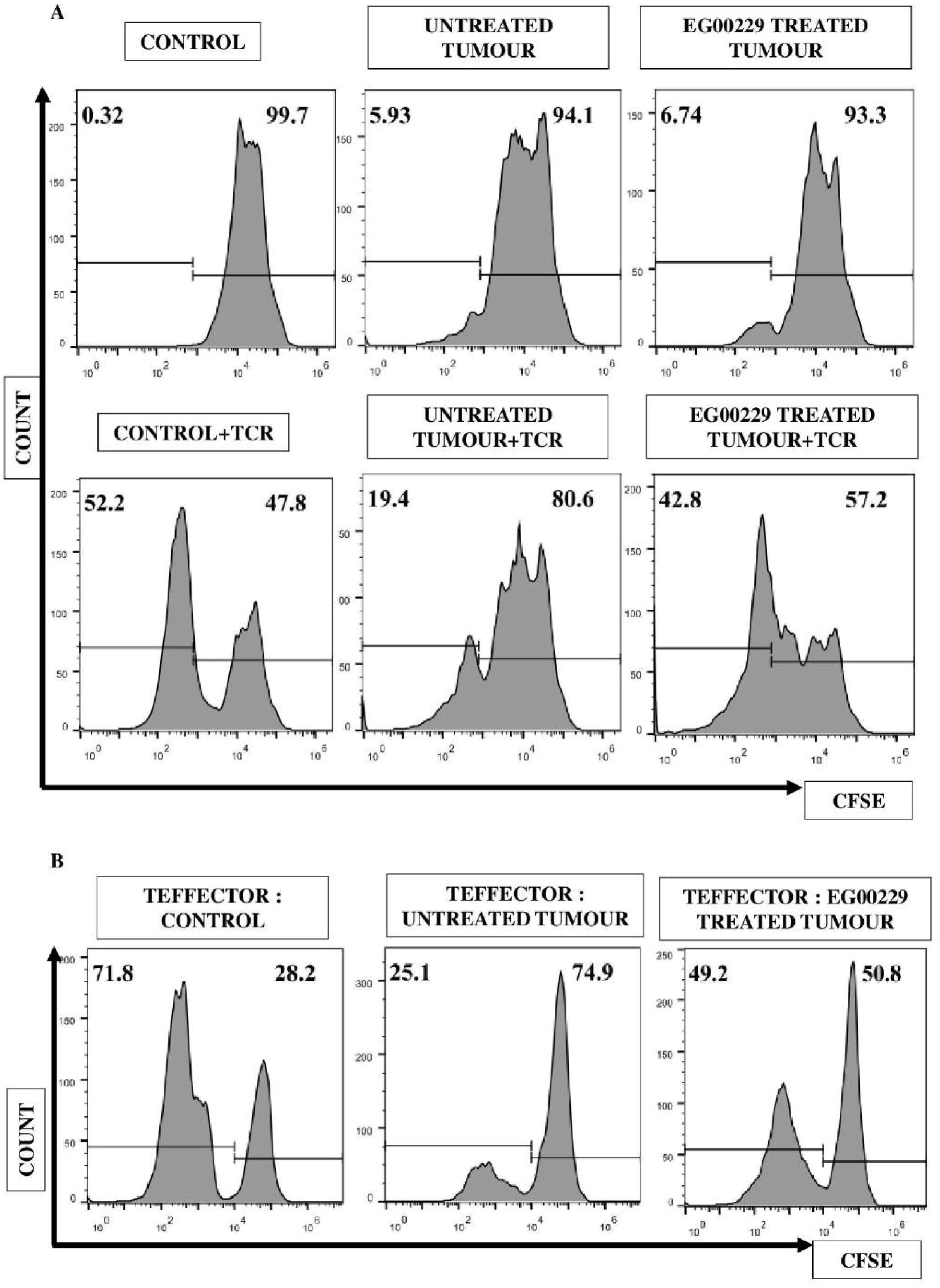
Increase in proliferation and reduction in suppressive activity of Tregs of purified splenic T cells isolated from B16-F10 subcutaneous tumour-bearing mice treated with Neuropilin-1 (NRP1) inhibitor. (A) Flowcytometric histogram plots depicting proliferation of purified splenic T cells isolated from B16-F10 subcutaneous tumour-bearing mice with or without treatment with Neuropillin-1 (NRP1) inhibitor. (B) Flowcytometric histogram plots depicting proliferation of effector T cells cultured with purified T cells isolated from B16-F10 subcutaneous tumour-bearing mice with or without NRP1 inhibitor treatment.

It was observed that TCR-activated splenic T cells isolated from EG-treated B16-F10 tumour-bearing mice showed increased proliferation compared to those isolated from untreated tumour-bearing mice (**Figure 7A and Supplementary figure S10A**). Furthermore, the absolute CD4^+^ NRP1^+^ Treg counts in murine spleens isolated from control, untreated tumour-bearing and EG-treated tumour-bearing C57BL/6 mice were evaluated (**Supplementary figure S6A and C**). It was found that the absolute count of CD4^+^ NRP1^+^ Tregs isolated from untreated tumour-bearing mice spleens was lower compared to EG-treated tumour-bearing mice.

Similarly, to evaluate the possible regulation of the suppressive activity of the T mentioned above cells, TCR-activated effector T cells were co-cultured with splenic T cells from untreated and EG-treated B16-F10 tumour-bearing mice. CD4^+^ splenic T cells isolated from EG-treated tumour-bearing mice showed lesser suppression of co-cultured effector T cell proliferation as compared to CD4^+^ splenic T cells isolated from untreated tumour-bearing mice (**Figure 7B and Supplementary figure S10B**).

Thus, EG treatment in B16-F10 tumour-bearing mice increased the proliferation of splenic T cells and reduced their immunosuppressive potential compared to T cells from untreated mice.

### Neuropillin-1 (NRP1) inhibitor treatment reduced B16-F10 lung metastasis and subcutaneous tumour volume in C57BL/6 mice, improving survival

NRP1 depletion, knockout and inhibition with inhibitors, monoclonal antibodies and nanobodies have shown remarkable anti-cancerous effects, including attenuated angiogenesis, tumorigenesis, tumour growth and metastasis and improved survivability (7,23,29–37,77). While various inhibitors and microRNAs have shown anti-cancer activity by inhibiting NRP1, EG00229 has been reported to be the most specific and characteristic inhibitor of NRP1 (28). It reportedly showed significant tumour-suppressive effects in cancers (38,39,86–88). Even though NRP1 has been reported as a potential target for melanoma and repurposed drugs have reportedly shown significant anti-tumour activity against melanoma (45,46), any possible anti-tumour activity of EG00229 on melanoma is yet to be studied. Accordingly, we evaluated the potential of the NRP1 inhibitor EG00229 trifluoroacetate (EG) to reduce the tumour growth and lung metastasis of B16-F10 melanoma tumours in C57BL/6 mice while improving their survival rates (**Figure 8**).

**Figure 8:**
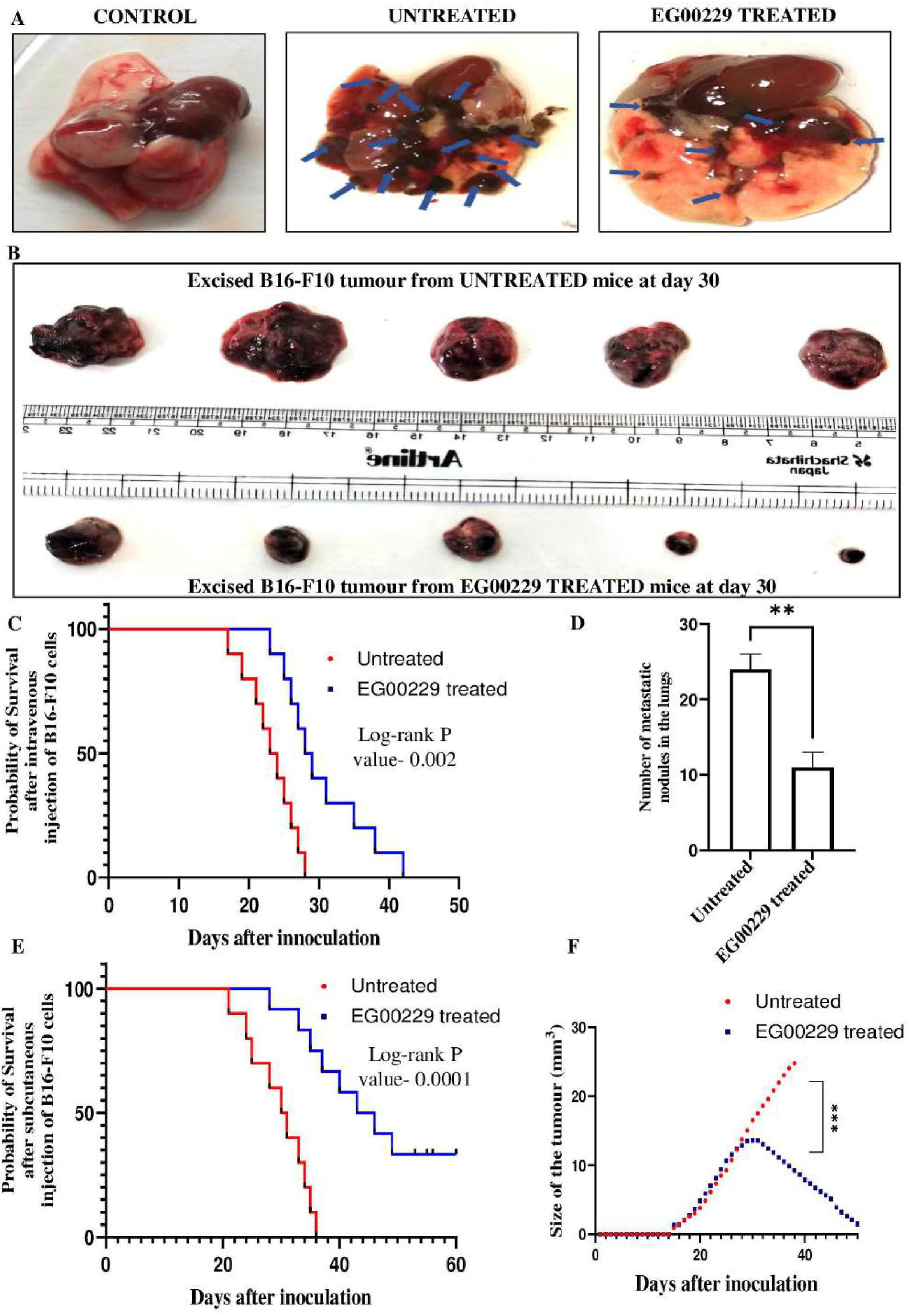
Neuropillin-1 (NRP1) inhibitor treatment alleviated B16-F10 lung metastasis and subcutaneous tumour volume in C57BL/6 mice and increased its survival. (A) Representative lung images of healthy control mice and lungs extracted from untreated and NRP1-treated C57BL/6 mice previously injected with B16-F10 cells intravenously. (B) Kaplan-Meier plot depicting the probability of survival of C57BL/6 mice with or without NRP1 inhibitor treatment after inoculation of 0.1X10^6^ B16-F10 cells intravenously. (C) Bar graph depicting the number of metastatic nodes in the lungs of control healthy mice and in lungs extracted from untreated and NRP1 treated C57BL/6 mice previously injected with B16-F10 cells intravenously. (D) Kaplan-Meier plot depicting the probability of survival of C57BL/6 mice with or without NRP1 inhibitor treatment after inoculation of 0.3X10^6^ B16-F10 cells subcutaneously. (E) XY plot depicting the volume of subcutaneous tumours in C57BL/6 mice with or without NRP1 inhibitor treatment after inoculation of 0.3X10^6^ B16-F10 cells subcutaneously.

Mice treated with EG showed fewer metastatic lung nodules from intravenous injection of B16-F10 melanoma cells than untreated mice (**Figure 8A and D**). Additionally, less subcutaneous tumour growth was observed in tumour-bearing mice treated with EG than in untreated mice (**Figure 8B and F and Supplementary figure S11**). According to log-rank P analysis, EG treatment improved survival rates in mice injected with B16-F10 cells subcutaneously or intravenously (**Figure 8C and E**).

Thus, treatment with EG reduced B16-F10 metastasis and tumour growth and improved the survivability rate in C57BL/6 mice.

## Discussion

A GLOBOCAN study identified melanoma as the deadliest skin cancer, posing a significant threat to public health and economic burden. Despite being preventable, melanoma is severe due to its high metastasis risk (89). Studies show that melanoma is immunogenic but has a poor prognosis in immunosuppressive conditions. Melanomas and other cancers recruit regulatory immune cells that inhibit effector T-cell function and express molecules that suppress activated immune cells (2,19). These studies led to the development of ICIs targeting PD-1/PD-L1 and CTLA-4, revolutionising cancer immunotherapy, especially against melanoma. New antibodies targeting LAG-3, TIM-3, VISTA, BRAF, mitogen-activated extracellular signal-regulated kinase (MEK), and T-cell immunoreceptor with Ig and ITIM domains (TIGIT) are currently in clinical trials alone or with anti-CTLA-4 and anti-PD-1/PD-L1. The success of these ICIs in melanoma has demonstrated their therapeutic potential. However, even with combination ICIs, around half of patients do not experience lasting benefits (21). IrAEs are also common, particularly with combination ICI therapy. Therefore, identifying molecular markers that regulate immune checkpoints and developing suitable immunotherapeutic regimens is crucial (11,23). Recently, NRP1 has been identified as a potential therapeutic target for cancer immunotherapy. NRP1 is reported to be upregulated in several cancers, promoting progression and poor prognosis (23,25,28,31,47,90). Inhibiting or depleting NRP1 reduces the immunosuppressive potential of intratumoral Tregs, angiogenesis, and tumour progression while enhancing other immunotherapies. Among the various inhibitors, monoclonal antibodies, and nanobodies targeting NRP1, EG00229 is the most specific inhibitor identified so far (7,27,32,37,67,91,92). Previous studies have determined that the NRP1 pathway is crucial for intratumoral Treg stability but unnecessary for maintaining immune homeostasis. This suggests that targeting NRP1 could be a viable strategy to limit tumour-induced tolerance without triggering autoimmunity, one of the significant IrAEs during ICI therapy (26). While EG00229’s anti-cancer potential has been demonstrated in different cancers, its role in melanoma and regulation of peripheral Tregs remains unknown. Accordingly, the potential immunotherapeutic role of EG00229 trifluoroacetate (EG) in regulating splenic peripheral Tregs during B16-F10 melanoma-driven immunosuppression was investigated. Also, its possible anti-tumorigenic effects in reducing B16-F10 melanoma tumour growth and metastasis and improving survival were evaluated.

While several studies have shown that Tregs, such as FOXP3^+^, NRP1^+^, and NKG2A^+^ Tregs, infiltrate tumours and are linked to poor prognosis in various cancers, their presence in peripheral sites during cancer-related immunosuppression remains underexplored (9,10,60). Additionally, while earlier reports have described B16-F10 melanoma tumour-mediated and B16-F10 cancer cell conditioned media (B16-F10-CS)-induced immunosuppression (50,63), the potential upregulation of Tregs during *in vitro,* B16-F10-CS-mediated immunosuppression has not yet been explored. It was observed that the frequency of FOXP3, NKG2A, and NRP1 positive Treg population increased in TCR-activated B16-F10-CS-treated peripheral splenic T cells compared to TCR-activated T cells alone (**Figure 1**). Meanwhile, upregulation of NKG2A and FOXP3 positive peripheral Tregs has been reported with head and neck squamous cell carcinoma (HNSCC) and gastric cancer (GC) cell conditioned media (61,62), respectively; the upregulation of NRP1^+^, NKG2A^+^, and FOXP3^+^ peripheral splenic Tregs in the presence of B16-F10-CS was observed for the first time. Additionally, the frequency of NRP1^+^ and NKG2A^+^ Tregs were upregulated only in B16-F10-CS-treated conditions compared to resting conditions (**Figure 1A, C and D**), while FOXP3^+^ Tregs frequency remained the same in both conditions, highlighting the importance of TCR-dependent FOXP3 upregulation, as reported earlier (93) (**Figure 1A and B**).

Furthermore, NRP1 is reported to be upregulated in various cancers and is essential for tumour cell survival, proliferation, and metastasis (23,25,26,31,44,91,94). Similarly, oncoprint from cBioPortal analysis of patient data from a pan-cancer study (ICGC/TCGA, Nature 2020) and a combined study of melanoma patients (Metastatic Melanoma (DFCI, Science 2015), Melanoma (MSK, NEJM 2014), and Metastatic Melanoma (UCLA, Cell 2016)) depicted that NRP1 gene expression was amplified in all cancer patients including melanoma (**Figure 2A**). Interestingly, it was observed from the oncoprint and the bar graph from cBioPortal analysis that the alteration frequency of the NRP1 gene was higher among melanoma cancer patients, suggesting NRP1 is a suitable therapeutic target for melanoma cancer as reported elsewhere (**Figure 2A and B**). Additionally, mRNA quantification showed elevated NRP1 gene expression in tumorigenic B16-F10 cells compared to non-tumorigenic NIH-3T3 cells (**Figure 2C**). Moreover, NRP1 was crucial for B16-F10 cell survival, as inhibition of NRP1 with EG induced apoptosis in B16-F10 cells at doses higher than 10 μM, but did not affect NIH-3T3 cells at the same dose (**Figure 2D, Supplementary figure S3**). The current findings validated earlier findings that NRP1 was indispensable for cancer cell survival and proliferation (23,67).

Earlier reports suggested that NRP1 is essential for both cancer cells and the stability and functioning of Tregs (26,90,95). Other NRP1 inhibitors have been shown to reduce the immunosuppressive potential of intratumoral Tregs (37). Although the NRP1 inhibitor EG is known for its anti-tumour activity in various cancers (41,47), its potential role in Tregs has remained unexplored until now. To the best of our knowledge, this study is the first to highlight the possible regulatory role of EG on peripheral Tregs. The findings revealed that NRP1 inhibition with EG in peripheral Tregs cultured in B16-F10-CS, *in vitro*, not only reduced their NRP1 expression but also decreased the expression of other immunosuppressive markers like NKG2A and FOXP3 (**Figure 3**), signifying the regulatory role of NRP1 in peripheral Tregs.

NRP1 inhibition also augmented T-cell effector responses in several cancers and facilitated anti-tumour responses (23,26). The current study examined the possible regulatory role of EG-mediated NRP1 inhibition in peripheral Tregs during B16-F10-CS *in vitro*. It was found that the EG-mediated NRP1 inhibition in TCR+B16-F10-CS-treated T cells not only led to their increment in effector cytokine secretion but also decreased their immunoregulatory cytokine secretion that is known to facilitate Treg-mediated immunosuppression otherwise (20,96) (**Figure 4A-F**). Although IL-17A has been reported to be an inflammatory cytokine, it has also been studied to promote Treg-mediated immunosuppression (97,98). In the current study, it has been observed that EG-mediated NRP1 inhibition in TCR+B16-F10-CS-treated splenic T cells downregulated their IL-17A secretion as compared to only TCR+B16-F10-CS treated ones, highlighting the role of NRP1 in IL-17A secretion and regulatory role in peripheral Tregs for the first time. NRP1 depletion has been reported to diminish FOXP3^+^ Treg proliferation (85). Similarly, TCR+B16-F10-CS+EG-treated CD4^+^ T cells also showed augmented proliferation compared to only TCR+B16-F10-CS-treated ones, which validated previous findings (**Figure 4G and Supplementary figure S7A**). NRP1 expression on Tregs is also known to enhance its immunosuppressive potential. Tregs facilitate immunosuppression during cancer by ameliorating effector T cell proliferation (99,100). In the current study, it was observed that TCR+B16-F10-CS+EG-treated peripheral splenic T cells augmented the proliferation of TCR-activated CFSE-labelled CD4^+^ effector T cells, thus highlighting that EG-mediated inhibition diminishes the immunosuppressive potential of peripheral Tregs, *in vitro* (**Figure 4H and Supplementary figure S7B**). It was also observed that TCR, B16-F10-CS and EG treatment did not alter the percentage of CD4^+^ T cells in each experimental condition (**Supplementary figure S5A and B**). Also, the absolute number of CD4^+^ NRP1^+^ Tregs decreased upon EG treatment in TCR and TCR+B16-F10-CS conditions compared to untreated TCR and TCR+B16-F10-CS conditions. This decrease in CD4^+^ NRP1^+^ Tregs in EG-treated conditions might have contributed to their diminished immunosuppressive potential observed here (**Figure 4H and Supplementary figure S7B**).

During cancer, NRP1 on Tregs facilitated immunosuppression by helping Tregs gather through VEGF co-receptor activity and maintaining their stability through Sema4a binding. This process promoted Treg survival and function by regulating the Akt-mTOR signalling and the PTEN-Akt-FoxO pathway. (24–26). It has been reported that Akt and PI3K phosphorylation is diminished during cancer conditions, and phosphorylated PTEN-mediated FOXP3 expression in Tregs is promoted (101). Accordingly, it was found that EG-mediated NRP1 inhibition in TCR+B16-F10-CS-treated peripheral Tregs showed elevated expression of phospho-Akt and PI3K as compared to only TCR+B16-F10-CS treated ones (**Figure 5A, F and G**). In addition, FOXP3 expression was also diminished in TCR+B16-F10-CS+EG-treated peripheral splenic T cells, thus validating previous findings (**Figure 2A and D**). Furthermore, it has also been reported that NRP1 on Tregs upregulated Smad2/3, ERK and P38 pathways through TGF-β and IL-10 mediated VEGFR and EGFR signalling (100,102). SAPK/JNK signalling is also reported to be downregulated in non-small-cell lung cancer cells (NSCLCC) through VEGFR signalling (103). Accordingly, it was found that EG-mediated NRP1 inhibition in TCR+B16-F10-CS-treated peripheral splenic T cells downregulated the expression of phospho-Smad2/3 and ERK as compared to only TCR+B16-F10-CS-treated ones (**Figure 5A, E and H**). However, no change in phosphorylated levels of P38 and SAPK/JNK was observed when compared between TCR+B16-F10-CS treated peripheral splenic T cells with or without EG treatment (**Figure 5A and Supplementary figure S8**). This indicates that NRP1 might not have a significant role in regulating SAPK/JNK and P38 signalling in peripheral Tregs, but further experimentations on NRP1-inhibited/depleted TILs isolated from B16-F10 tumours must be conducted to validate this hypothesis. Moreover, while the significance of STAT signalling in different cancers and regulation of NRP1-mediated immunosuppression of Tregs through STAT signalling has been extensively studied, the regulatory role of NRP1 in STAT signalling in peripheral Tregs is understudied (80,104–108). Earlier findings suggested that upregulation in phosphorylated STAT3 and STAT5 expression in Tregs is mediated through IL-10 and TGF-β signalling and is reported to augment FOXP3 expression and immunosuppressive potential of Tregs (109,110). Similarly, in the current study, it was observed that NRP1 inhibition in TCR+B16-F10-CS-treated peripheral splenic T cells downregulated the expression of phosphorylated STAT3, STAT5 and STAT1 as compared to only TCR+B16-F10-CS treated ones (**Figure 5A-D**). STAT1 is typically regarded as a tumour suppressor but can exhibit complex functions in tumour cells and the immune system. Inappropriate activation of STAT1 has been observed in various malignancies, including breast cancer, head and neck squamous carcinoma, melanoma, lymphoma, and leukaemia, suggesting that it may promote malignant transformation under certain conditions. Additionally, STAT1 is essential for inducing the expression of IDO, which enhances tryptophan catabolism and blocks T-cell activation (105). Therefore, it might indicate that EG-mediated NRP1 inhibition in TCR+B16-F10-CS-treated peripheral splenic Tregs downregulated STAT, Smad2/3 and ERK pathways that are reported to aid in immunosuppression in Tregs. It also upregulated AKT/PI3K pathways that are reported to destabilise FOXP3 expressing Tregs (111).

NRP1 also inhibited the immunosuppressive Treg population in intratumoral regions in different cancers (64,65,71–75). The current study depicted its possible role in regulating the expression of immunosuppressive markers in peripheral splenic Tregs. Peripheral splenic T cells isolated from EG-treated B16-F10 tumour-bearing mice not only showed alleviated expression of different Treg markers like NRP1, NKG2A, FOXP3, PD-1, PD-L1 and CTLA-4 but also Th_2_ marker GATA-3 (**Figure 6A-G and Supplementary figure S9A-G**). Therefore, it was found that NRP1 inhibition downregulated the expression of Treg mar in peripheral Tregs as well. Moreover, although T-bet is known to be a marker for Tregs, it is reported to facilitate Th_1_ responses in Tregs (112). Interestingly, peripheral splenic T cells isolated from EG-treated B16-F10 tumour-bearing mice showed elevated expression of T-bet (**Figure 6H and Supplementary figure S4H**). Therefore, it was observed that NRP1 inhibition aided in promoting Th_1_ responses in Tregs that might help in anti-tumour activity since prolonged exposure to Th_1_ conditions was reported to be essential for sustained tumour-specific cytotoxic T lymphocyte activity (113).

Additionally, it was observed that peripheral splenic T cells isolated from EG-treated B16-F10 tumour-bearing mice showed elevated T cell proliferation (**Figure 7A and Supplementary figure S10A)**. It was also found that while there was a significant increase in CD4^+^ T cells in the spleens of tumour-bearing mice compared to control mice, there was no change in CD4^+^ T cell count in the spleens of untreated and EG-treated tumour-bearing mice (**Supplementary figure S5A and C**). However, the total number of CD4^+^ NRP1^+^ Tregs was lower in the spleens of EG-treated tumour-bearing mice than those untreated (**Supplementary figure S6A and C**). These cells isolated from spleens of EG-treated tumour-bearing mice containing a lower number of CD4^+^ NRP1^+^ Tregs also enhanced the proliferation of TCR-activated CFSE-labelled effector T cells (**Figure 7B and Supplementary figure S7B**). Thus, EG treatment in B16-F10 tumour-bearing mice increased peripheral splenic T cell proliferation and reduced Treg suppressive activity, unlike splenic T cells from untreated B16-F10 tumour-bearing mice. Overall, EG treatment augmented effector T cell proliferation and mitigated Treg suppressive activity in peripheral Tregs both *in vitro* and *in vivo* (**Figure 4G, 4H, 7A and 7B**).

NRP1 has been reported as a potential target for melanoma (44,65,114). Repurposed drugs like gliclazide and entrectenib have shown significant anti-tumour activity in metastatic melanoma by preventing the PDGF-C/NRP1 interaction (43). However, the possible anti-tumour activity of EG on melanoma has yet to be deciphered. It was observed that EG treatment diminished B16-F10 lung metastasis in syngenic C57BL/6 mice and reduced the formation of metastatic lung nodules (Figures **8A and D**). EG treatment also regressed the B16-F10-melanoma tumour volume (**Figure 8B and F and Supplementary figure S11**). Additionally, EG treatment enhanced the survivability of C57BL/6 mice injected with B16-F10 melanoma cells either intravenously or subcutaneously (**Figure 8C and E**). This may be due to tumour cell death from EG treatment, as our findings show, which can reduce peripheral Treg-mediated immunosuppression or a combination of both effects (**Figure 2D, 4, 6 and 7**).

The current study demonstrated for the first time that EG treatment might regulate peripheral Treg-mediated immunosuppression and inhibit B16-F10 melanoma tumour growth and metastasis while improving survival. Nonetheless, the possible regulation of EG-mediated NRP1 inhibition in intratumoral Tregs and possible anti-tumoral activity of EG against melanoma cancer in humans is to be conducted for a better evaluation of EG as a potential anti-cancer immunotherapeutic strategy. Moreover, as reported earlier (42), EG might be formulated with better delivery molecules to improve their dosage and efficient drug concentration. Furthermore, EG, in combination with other ICIs, must be examined for possible enhanced efficacy without causing any deleterious side effects, as seen for other ICIs.

## Conclusion

This study showed for the first time that EG treatment may control peripheral Treg-mediated immune suppression, slow the growth and metastasis of B16-F10 melanoma tumours, and help improve survival. This might highlight the potential of NRP1 as a promising anti-cancerous immunotherapeutic target, alone or in combination with the current standard therapeutic regimen.

## Supporting information

"C:\Users\Somlata Khamaru\Downloads\Supplementary figures with legend-compressed.docx"

## Ethics statement

The animal study was reviewed and approved by the Institutional Animal Ethics Committee, NISER (1634/GO/ReBi/S/12/CPSCEA) under the affiliation of the Committee for the Control and Supervision of Experiments on Animals (CCSEA) of India.

## Authors contribution

**Somlata Khamaru**: Contributed majorly to conceptualisation, design, all experimentation, data acquisition, data analysis, manuscript writing and editing. **Kshyama Subhadarsini Tung**: Contributed to manuscript writing, reviewing and editing. **Subhasis Chattopadhyay**: Contributed to conceiving, conceptualising, design of the study, fund acquisition, manuscript review and editing.

## Acknowledgement

The authors are thankful towards NISER Flowcytometry and the Animal House facility for providing pivotal support to conduct the study.

## Data availability

The original contributions presented in the study are included in the article/Supplementary Material. Further inquiries can be directed to the corresponding authors.

---

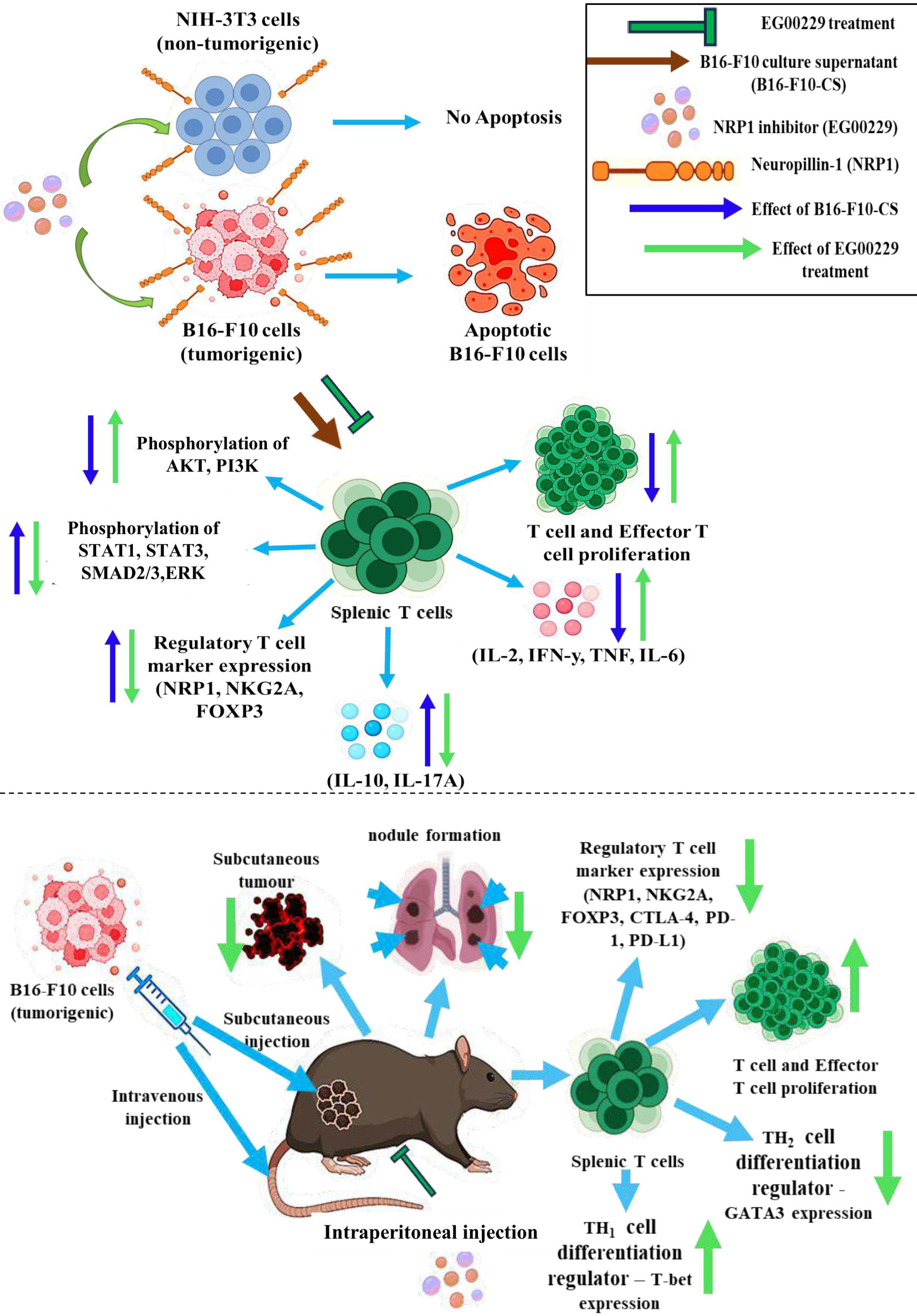

