## Supplementary material for "Targeting Neuropilin-1 to Enhance Immunotherapy in Melanoma: Reducing Peripheral Treg-Mediated Immunosuppression and Tumour Progression": "C:\Users\Somlata Khamaru\Downloads\Supplementary figures with legend-compressed.docx"

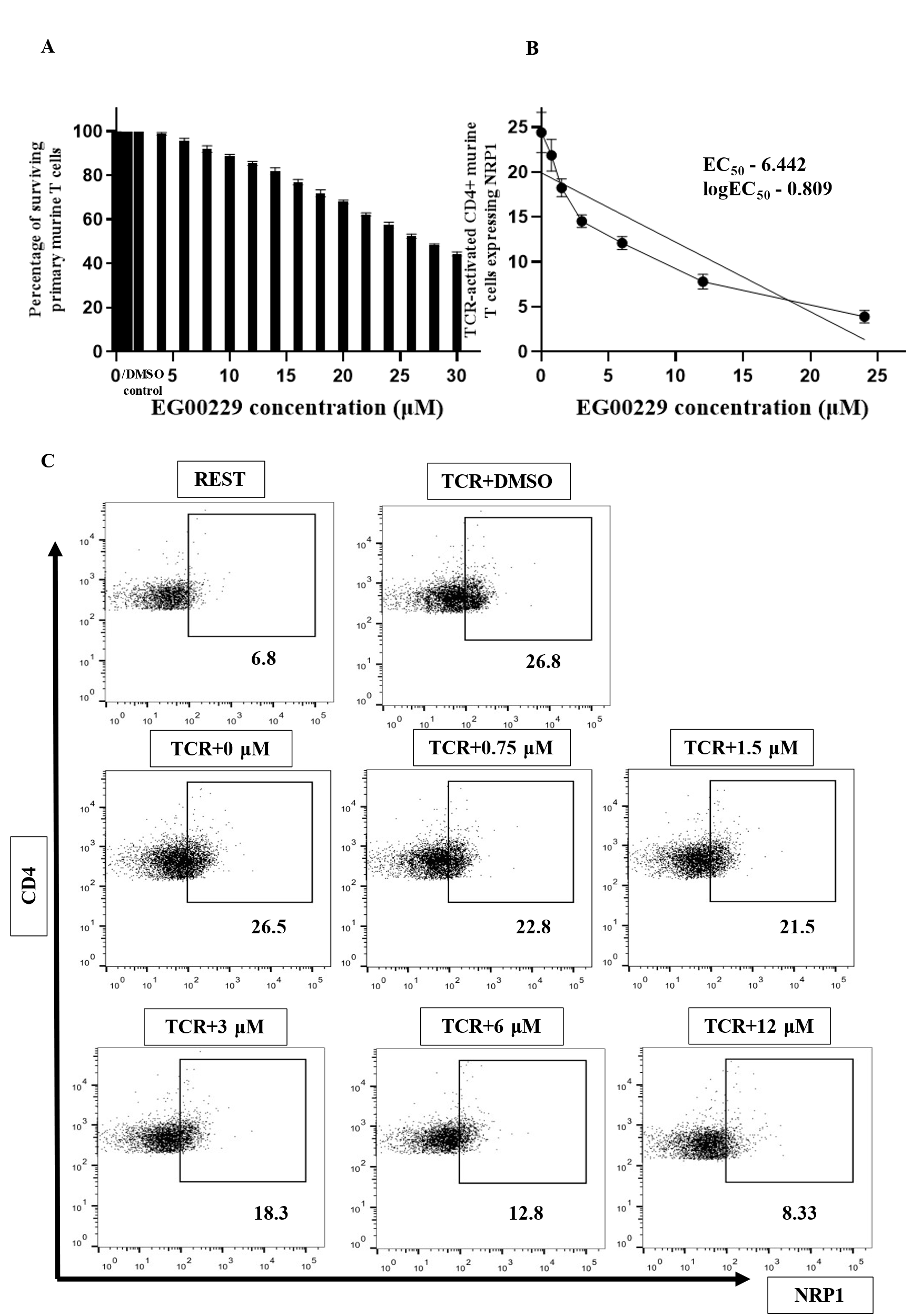

**Supplementary figure S1: Estimation of non-toxic dose and EC_50_ value of EG00229 trifluoroacetate (EG) on purified murine splenic T cells.** (A) The bar graph from the trypan blue exclusion test represents the percentage of surviving purified murine splenic T cells cultured in the presence of DMSO and NRP1 inhibitor EG at different concentrations. (B) Representative XY plot depicting the percentage of TCR-activated CD4+ NRP1+ purified murine splenic T cells upon treatment with NRP1 inhibitor EG. (C) Representative flow cytometric dot plots depicting the percentage of TCR-activated CD4+ NRP1+ purified murine splenic T cells upon treatment with NRP1 inhibitor EG. Representative bar and XY graphs are of three independent experiments.

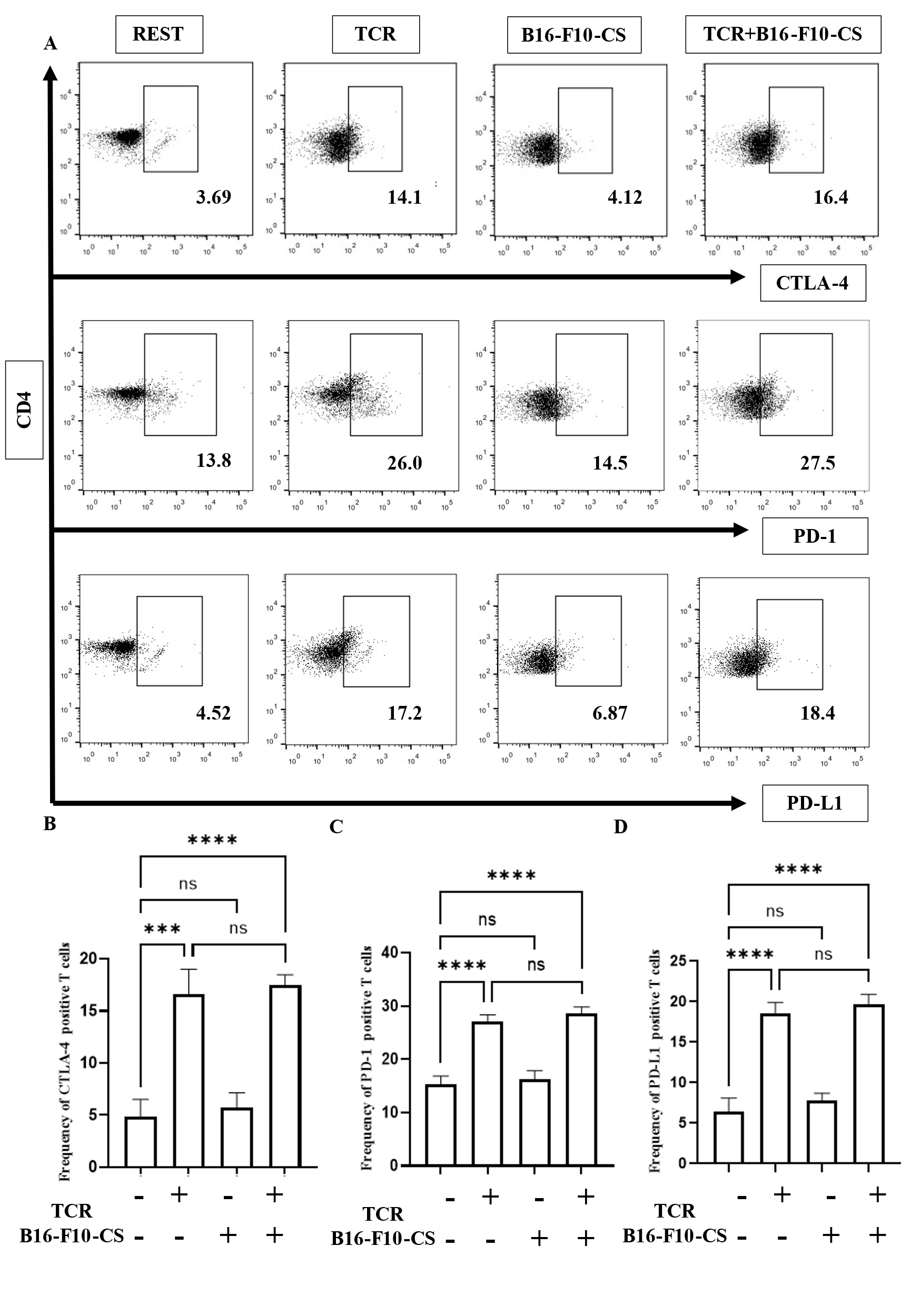

**Supplementary figure S2: Frequency of suppressor T cells during B16-F10 cell culture supernatant (B16-F10-CS) treatment, *in vitro*.** (A) Representative flowcytometric dot plots showing alterations in the frequency of CTLA-4, PD-1 and PD-L1 positive purified splenic T cell population with or without anti-CD3/CD28 antibodies and B16-F10-CS treatment. Bar graph representation of alteration in frequency of (B) CTLA-4, (C) PD-1 and (D) PD-L1 positive purified splenic T cell population with or without anti-CD3/CD28 antibodies and B16-F10-CS treatment. Representative bar diagrams are of three independent experiments. ns, non-significant; * p < 0.05; ** p < 0.01; *** p < 0.001; **** p < 0.0001.

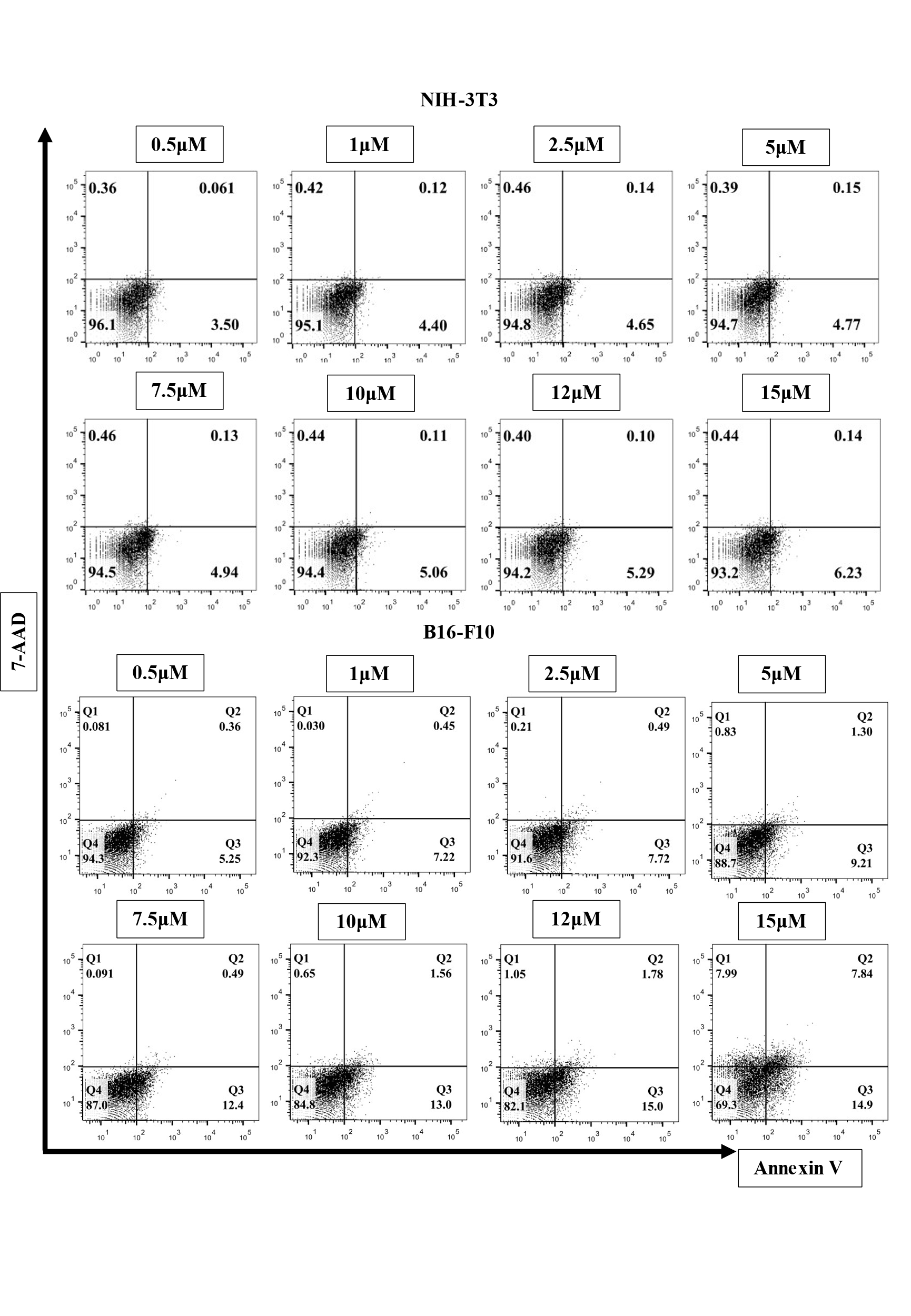
**Supplementary figure S3: Inhibition of Neuropillin-1 (NRP1) induced apoptosis in tumorigenic B16-F10 cells but not in non-tumorigenic NIH-3T3 cells.** Representative flowcytometric dot plots depicting apoptosis in tumorigenic B16-F10 and non-tumorigenic NIH-3T3 cells upon treatment with different concentrations of NRP1 inhibitor EG00229, which is determined by Annexin & 7-AAD dual staining.

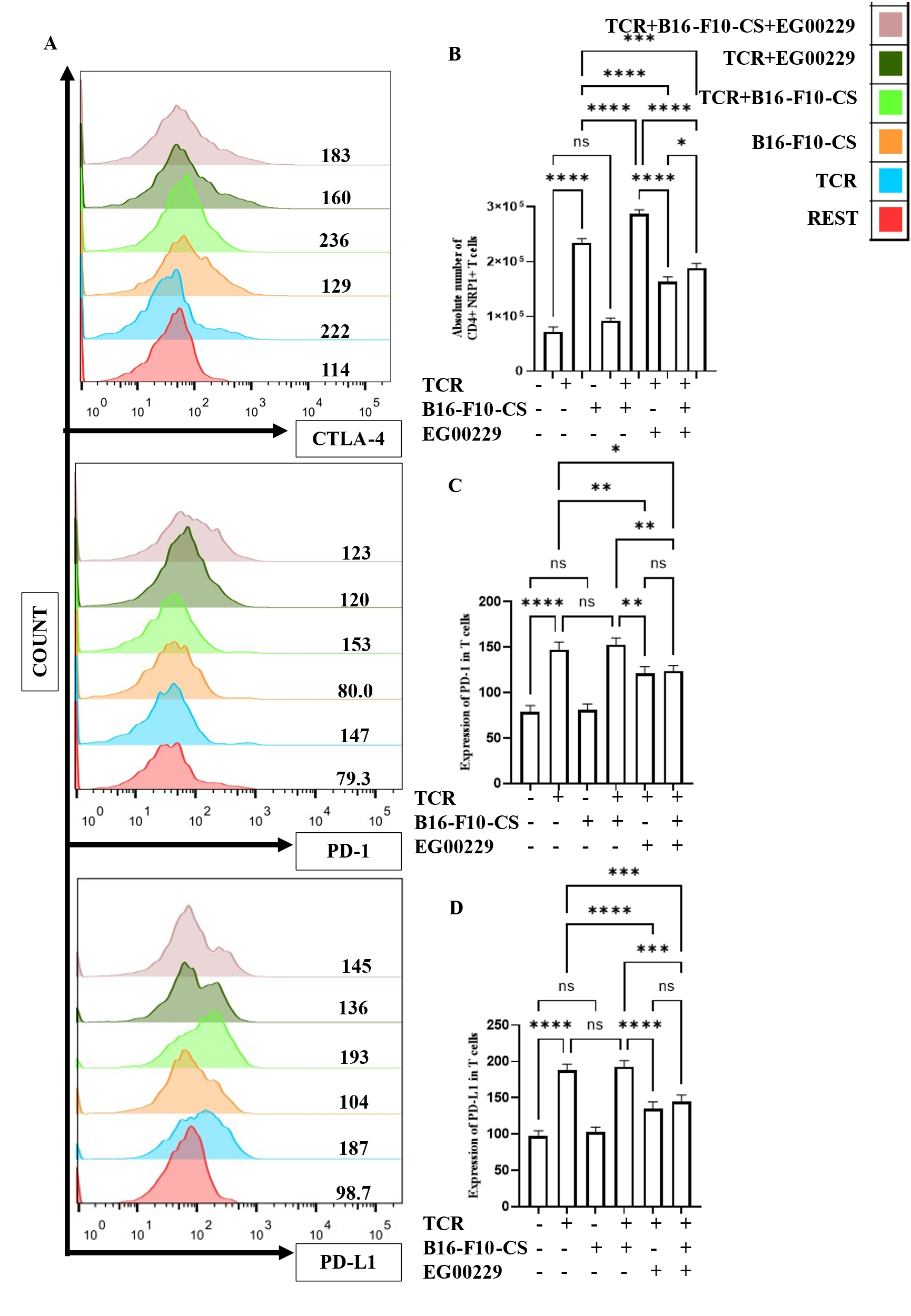
**Supplementary figure S4: Expression of suppressor T cell markers in Neuropillin-1 (NRP1) inhibited splenic T cells treated with B16-F10 cell culture supernatant (B16-F10-CS), *in vitro*.** (A) Representative flow cytometric histogram plots depicting alterations in expression of CTLA-4, PD-1 and PD-L1 in anti-CD3/CD28 antibodies and B16-F10-CS treated purified splenic T cell with or without NRP1 inhibition. Bar graph representation of alteration in expression of (B) CTLA-4, (C) PD-1 and (D) PD-L1 in anti-CD3/CD28 antibodies and B16-F10-CS treated purified splenic T cell with or without NRP1 inhibition. Representative bar diagrams are of three independent experiments. ns, non-significant; * p < 0.05; ** p < 0.01; *** p < 0.001; **** p < 0.0001.

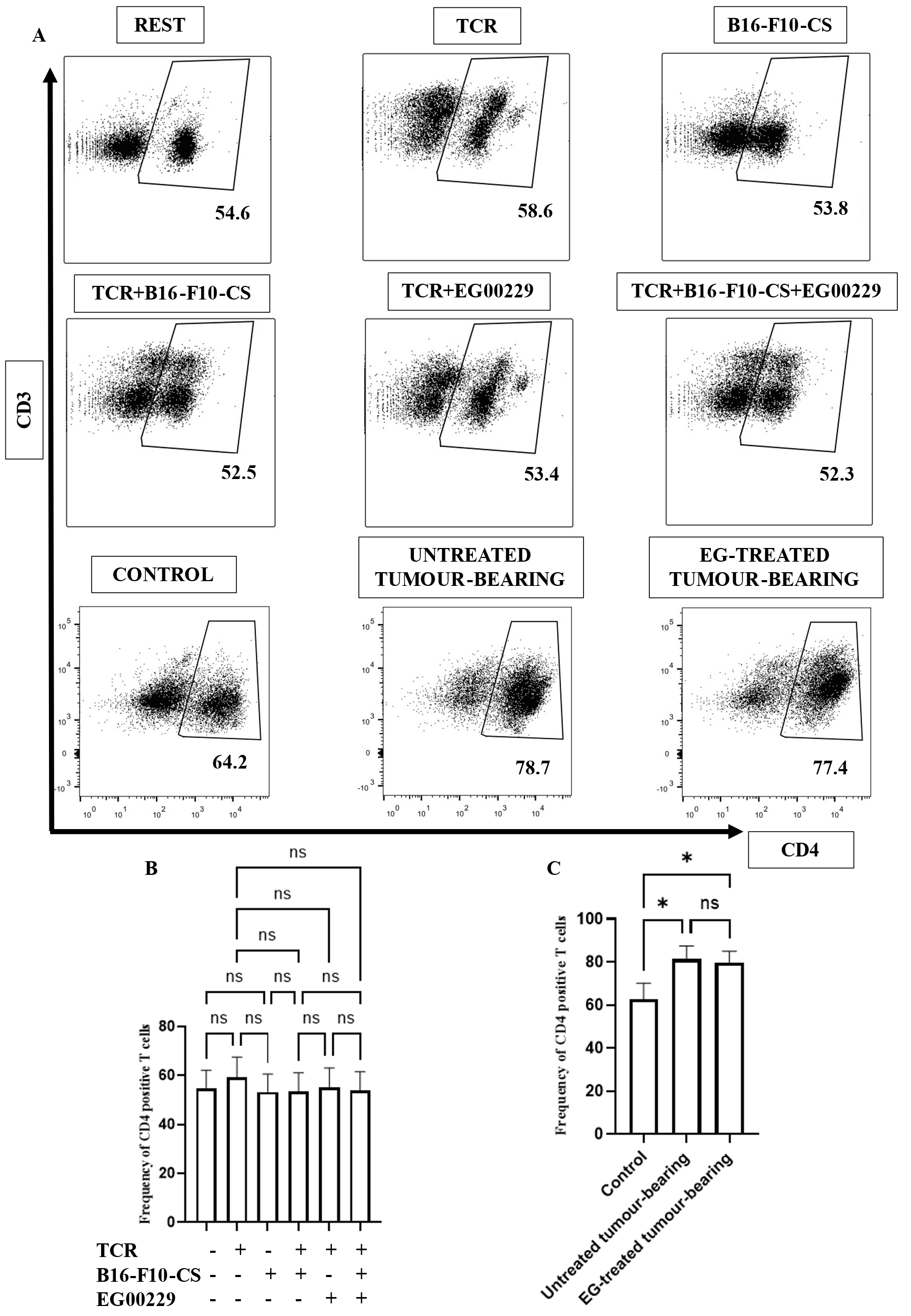

**Supplementary figure S5: Frequency of CD4+ purified murine splenic T cell population isolated from EG-treated or untreated tumour-bearing mice and control mice with or without *in vitro* EG and B16-F10-CS treatment.** (A) Representative flow cytometric dot plots depicting the frequency of CD4+ purified murine splenic T cell population isolated from EG-treated or untreated tumour-bearing mice and control mice with or without in vitro EG and B16-F10-CS treatment. Bar graph representation of the frequency of CD4+ purified murine splenic T cell population isolated from (B) control mice with or without in vitro EG and B16-F10-CS treatment, and (C) EG-treated or untreated tumour-bearing C57BL/6 mice. Representative bar diagrams are of three independent experiments. ns, non-significant; * p < 0.05; ** p < 0.01; *** p < 0.001; **** p < 0.0001.

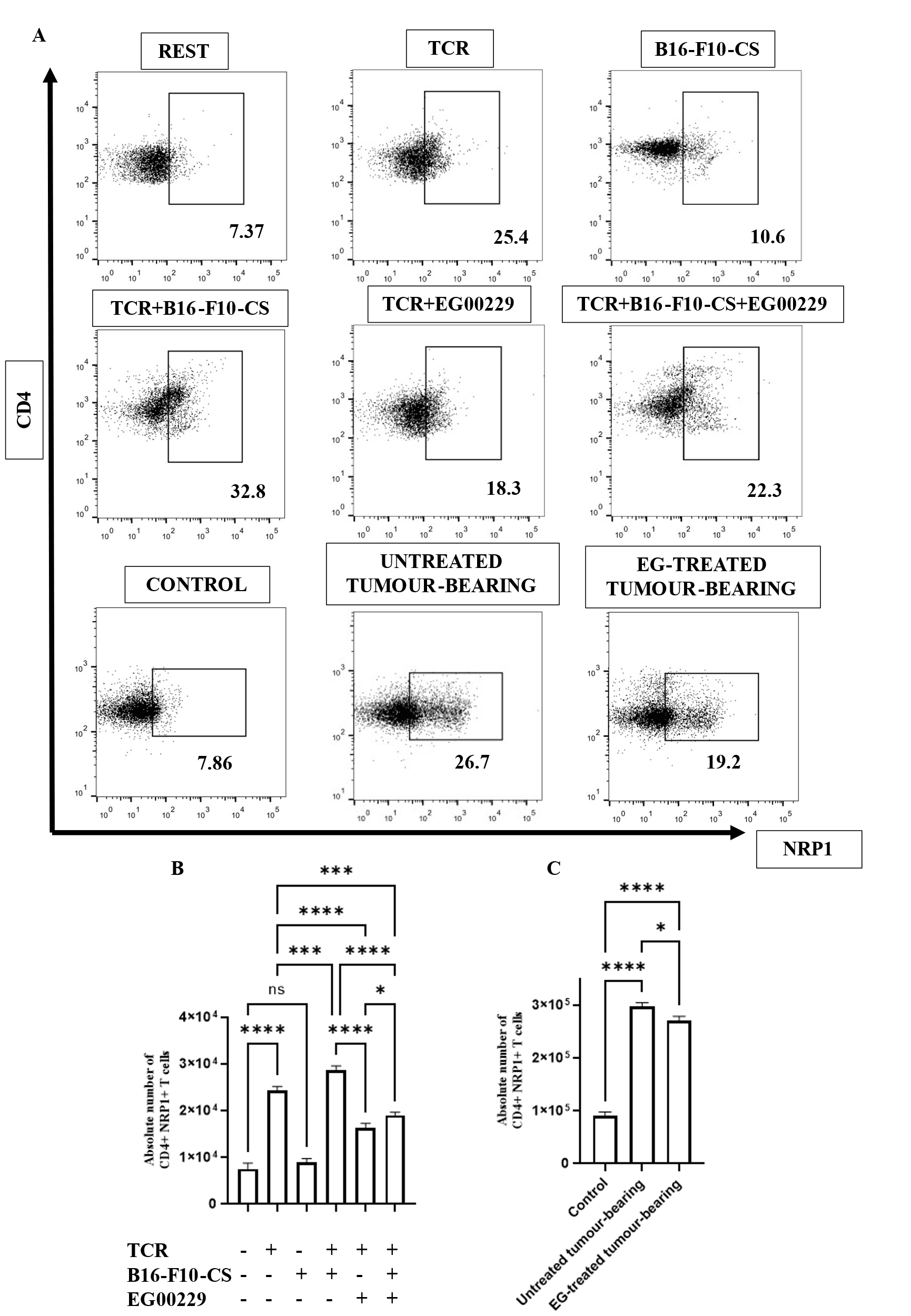
**Supplementary figure S6: Estimation of absolute number of CD4+ NRP1+ regulatory T cell (Treg) population isolated from EG-treated or untreated tumour-bearing mice and control mice with or without in vitro EG and B16-F10-CS treatment.** (A) Representative flow cytometric dot plots depict the frequency of CD4+ NRP1+ regulatory T cell (Treg) population isolated from EG-treated or untreated tumour-bearing mice and control mice with or without vitro EG and B16-F10-CS treatment. Bar graph representation of the estimated absolute number of CD4+ NRP1+ regulatory T cell (Treg) isolated from (B) control mice with or without in vitro EG and B16-F10-CS treatment, and (C) EG-treated or untreated tumour-bearing C57BL/6 mice. Representative bar diagrams are of three independent experiments. ns, non-significant; * p < 0.05; ** p < 0.01; *** p < 0.001; **** p < 0.0001.

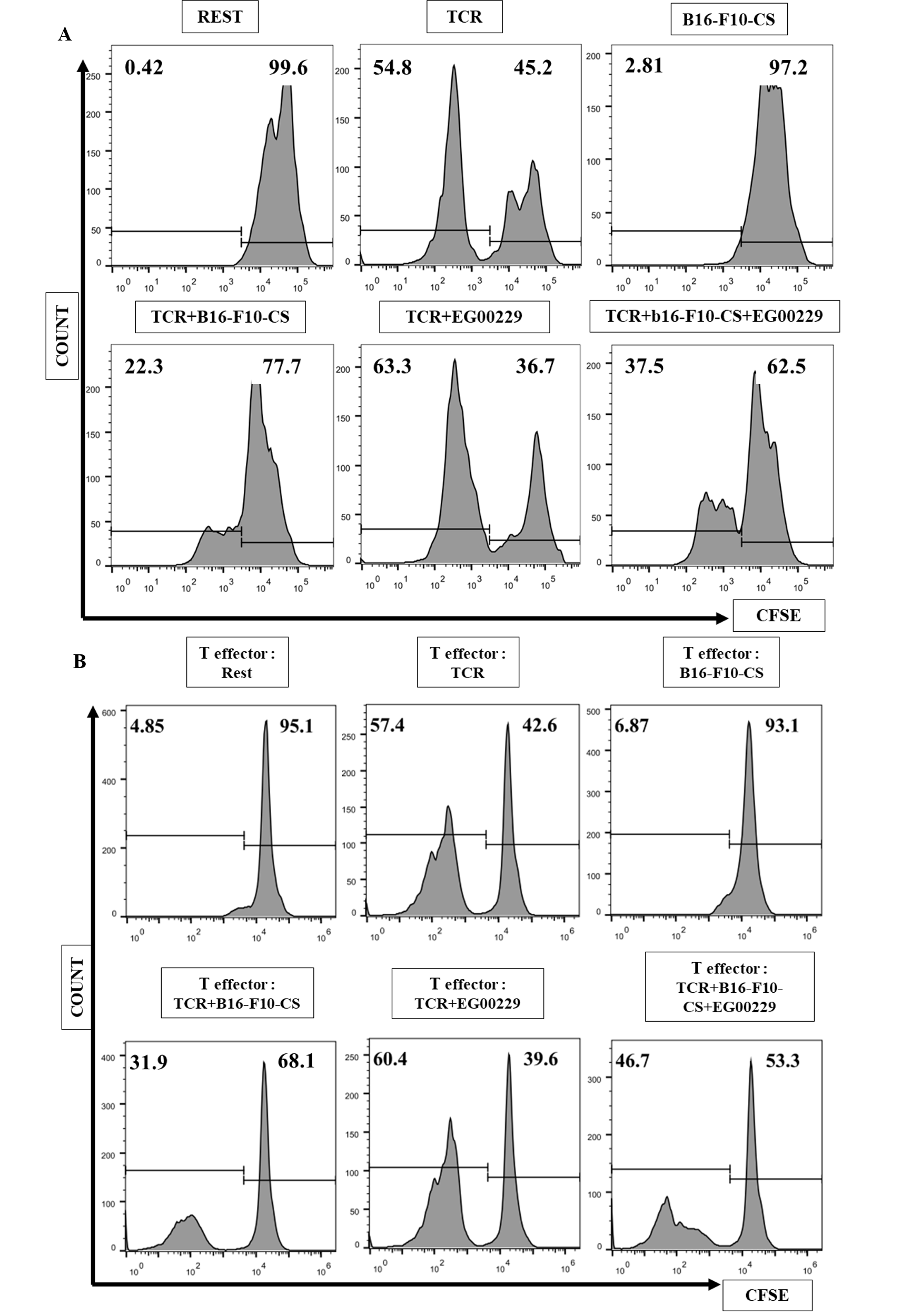
**Supplementary figure S7: Inhibition of Neuropillin-1 (NRP1) elevated T cell proliferation in splenic T cells treated with B16-F10 cell culture supernatant (B16-F10-CS) treatment, *in vitro*.** (A) Representative flow cytometric histogram plot depicting proliferation of T cells via CFSE proliferation assay upon treatment with anti-CD3/CD28 antibodies and B16-F10-CS with or without NRP1 inhibition. (B) Representative flow cytometric histogram plot depicting proliferation of effector T cells cultured in purified splenic T cells treated with anti-CD3/CD28 antibodies and B16-F10-CS with or without NRP1 inhibition.

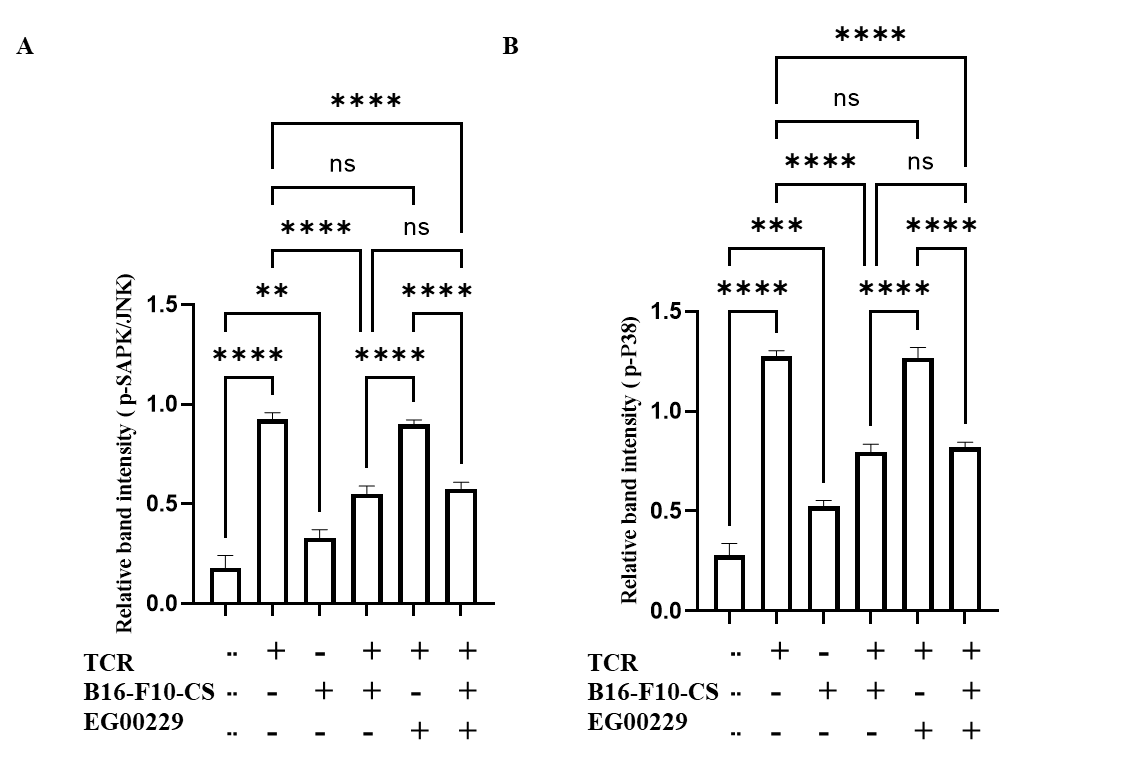

**Supplementary figure S8: Inhibition of Neuropillin-1 (NRP1) did not alter SAPK/JNK and P38 MAPK pathways in splenic T cells treated with B16-F10 cell culture supernatant (B16-F10-CS) treatment, *in vitro.*** Representative bar graphs are of three independent experiments, showing relative expression levels of (A) p-SAPK/JNK and (B) p-P38 when compared with the expression of total SAPK/JNK and P38, respectively. ns, non-significant; * p < 0.05; ** p < 0.01; *** p < 0.001; **** p < 0.0001.

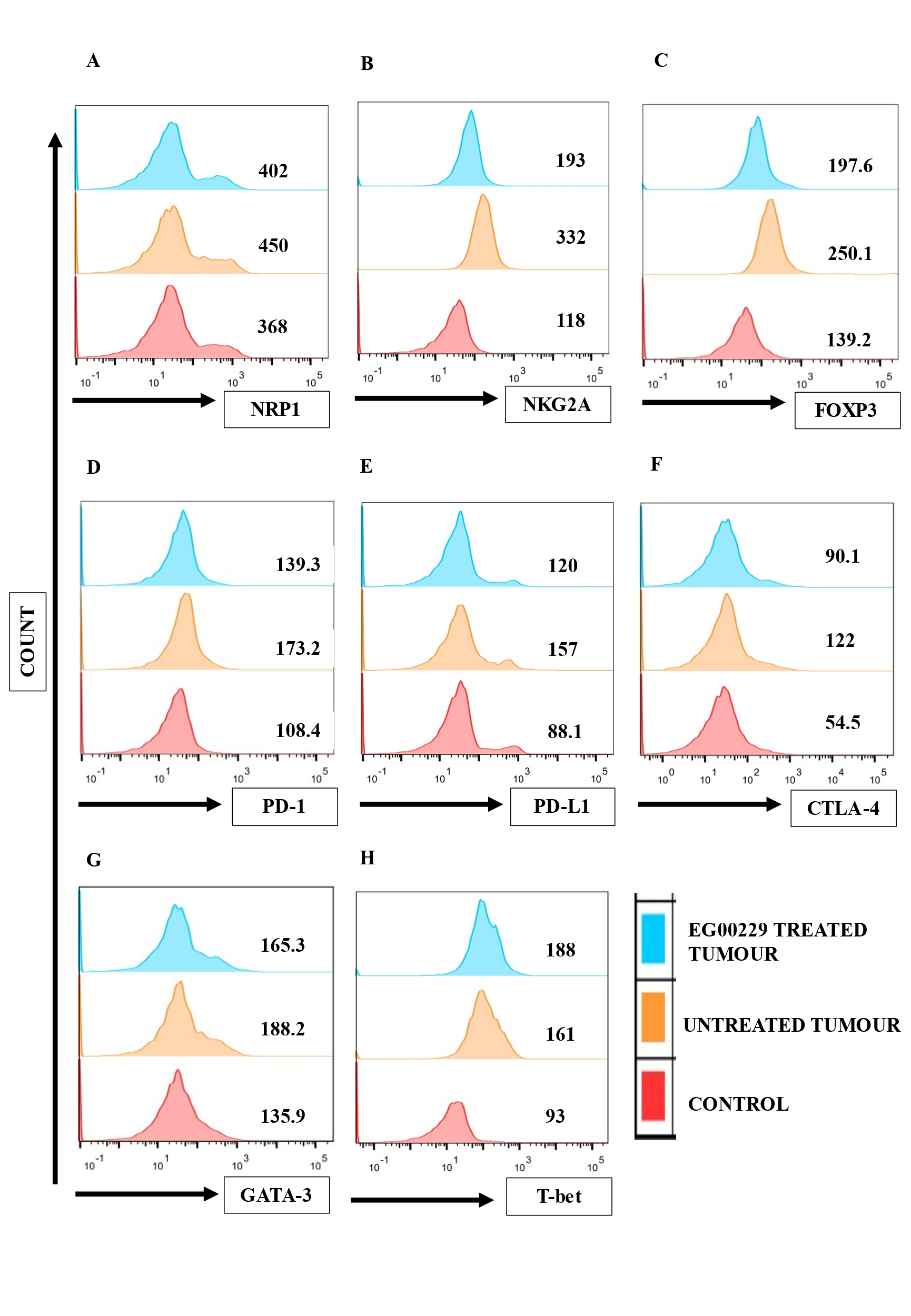

**Supplementary figure S9: Expression of suppressor T cell markers alleviated in B16-F10 subcutaneous tumour bearing mice treated with Neuropillin-1 (NRP1) inhibitor.** Representative flowcytometric histogram plots depicting the alteration in expression of (A) NRP1, (B) NKG2A, (C) FOXP3, (D) CTLA-4, (E) PD-1, (F) PD-L1, (G) T-bet and (H) GATA-3 in purified splenic T cells isolated from control healthy and B16-F10 subcutaneous tumour bearing C57BL/6 mice with or without NRP1 inhibitor treatment.

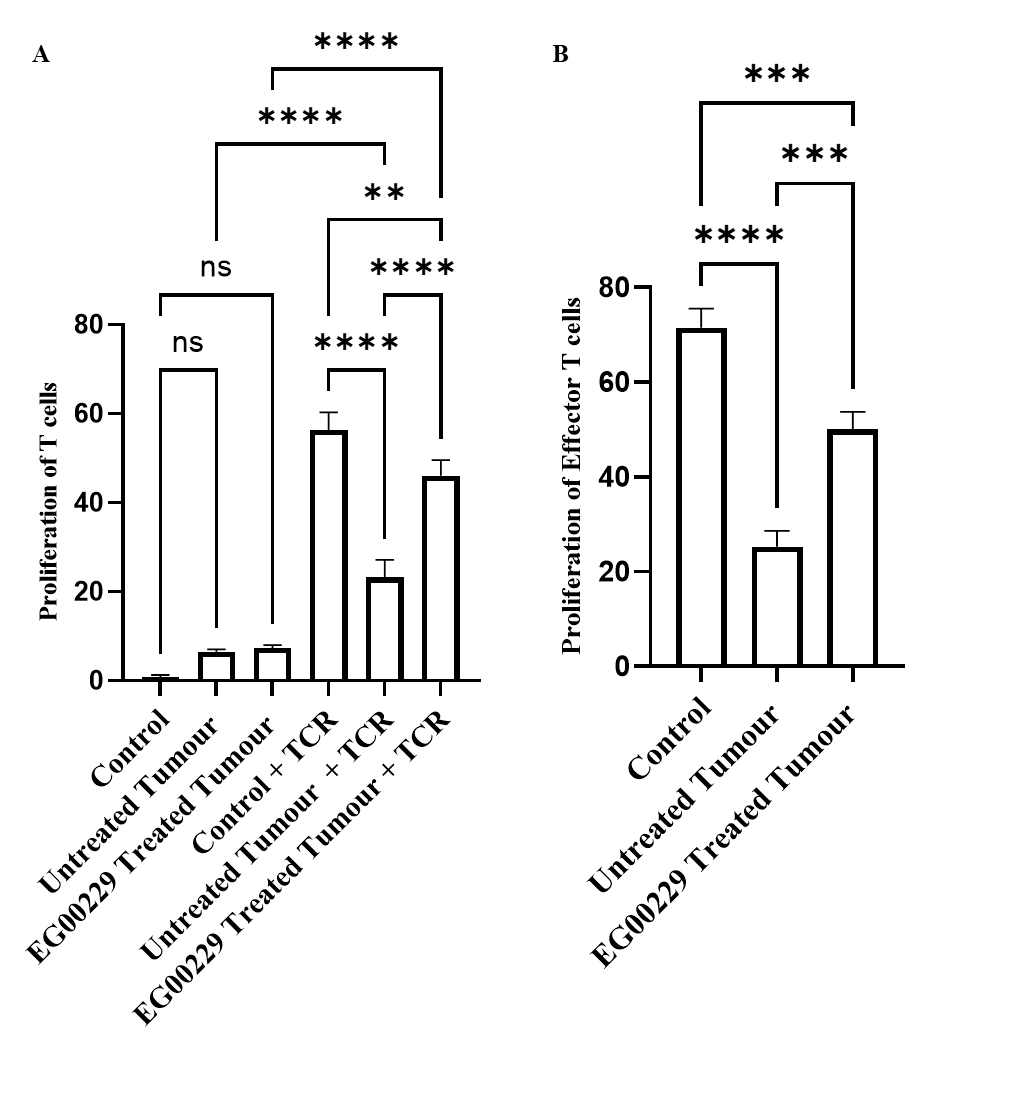

**Supplementary figure S10: Elevation in the proliferation of purified splenic T cells isolated from B16-F10 subcutaneous tumour-bearing mice treated with Neuropillin-1 (NRP1) inhibitor.** (A) Bar graph depicting proliferation of purified splenic T cells isolated from B16-F10 subcutaneous tumour-bearing mice with or without treatment with Neuropillin-1 (NRP1) inhibitor. (B) Bar graph depicting proliferation of effector T cells cultured with purified T cells isolated from B16-F10 subcutaneous tumour-bearing mice with or without NRP1 inhibitor treatment. Representative bar diagrams are of three independent experiments. ns, non-significant; * p < 0.05; ** p < 0.01; *** p < 0.001; **** p < 0.0001.

**Supplementary figure S11: Neuropillin-1 (NRP1) inhibitor treatment alleviated B16-F10 subcutaneous tumour volume in C57BL/6 mice.** Representative tumour images at day 18, 21, 24, 30 and 36 on C57BL/6 mice with or without NRP1 inhibitor treatment after inoculation of 0.3X10^6^ B16-F10 cells subcutaneously.

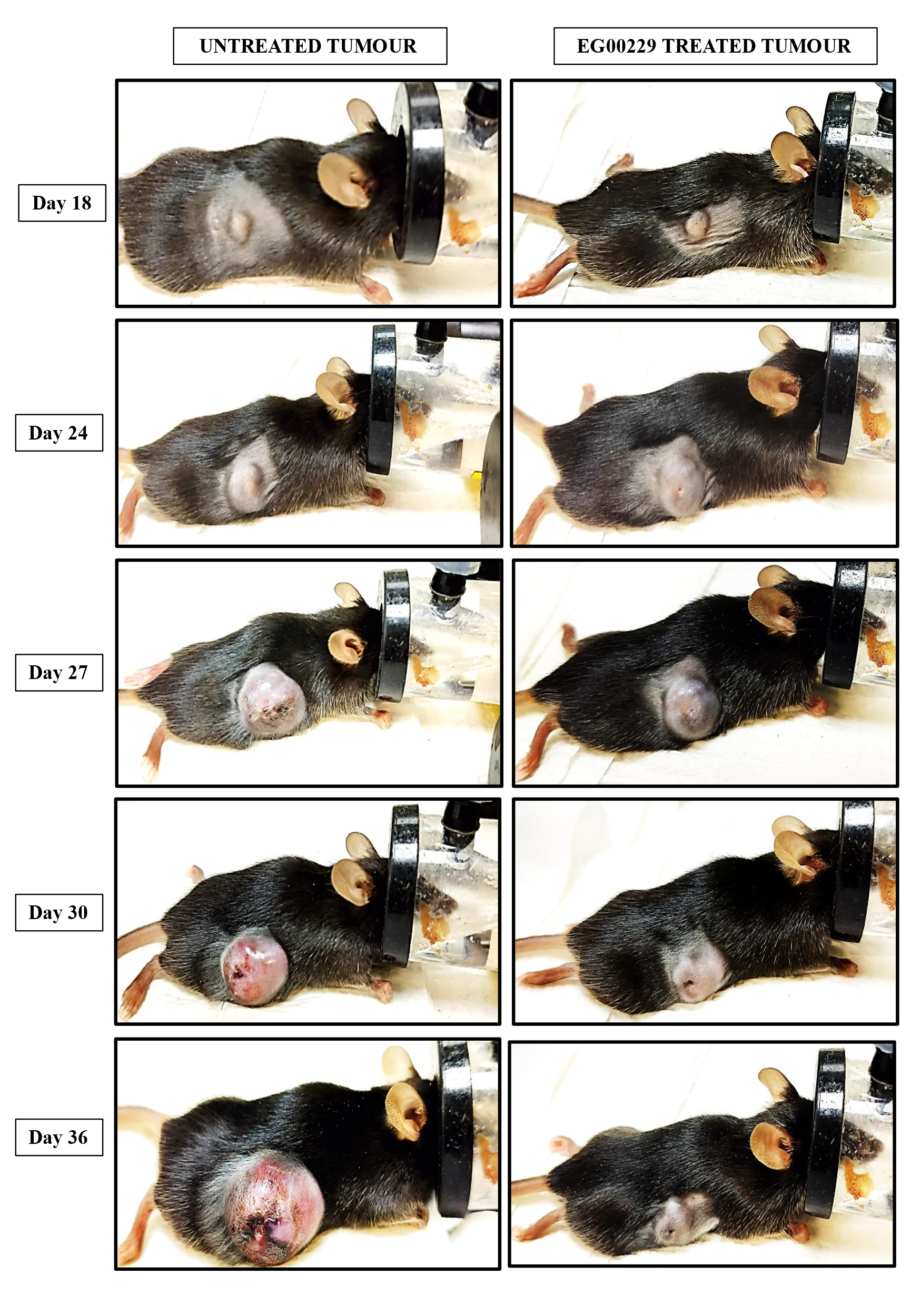
